# Encoding of visual stimuli and behavioral goals in distinct anatomical areas of monkey ventrolateral prefrontal cortex

**DOI:** 10.1101/2025.01.30.635437

**Authors:** C. Basile, M. Gerbella, A. Gravante, A. Lapadula, F. Rodà, L. Simone, L. Fogassi, S. Rozzi

## Abstract

The lateral prefrontal cortex has been classically defined as an associative region involved in the so-called executive functions, such as guiding behavior based on abstract rules and mnemonic information. However, most neurophysiological studies on monkeys did not address the issue of whether distinct anatomical sectors of lateral prefrontal cortex play different functional roles. The main aim of this work is to study functional properties of neurons recorded from a large part of ventrolateral prefrontal cortex (VLPF) of two monkeys performing passive visual tasks and a visuo-motor task, and to map them on the anatomical areas defined on the basis of our recent parcellations.

Our results show that distinct VLPF areas differently contribute to visual processing and action organization along the caudo-rostral axis. In particular, the processing of visual stimuli, independent of whether passively presented or exploited for guiding behavior, primarily involves posterior VLPF areas (especially caudal area 12r), while the elaboration of visual and contextual information for action organization mainly involves intermediate VLPF areas (especially middle 46v). In this latter sector, visual stimuli/instructions appear to be encoded in a pragmatic format, that is in terms of the associated behavioral outcome. Finally, more anterior areas are characterized by a low responsiveness to the employed tasks.

Altogether, our findings indicate that posterior VLPF areas represent the first processing stage of visual input, intermediate areas primarily contribute to the selection and planning of contextually appropriate behaviors, while rostral areas could be involved in more complex abstract processes.

## Introduction

The ability to adapt goal-directed behavior to a continuously changing context is fundamental for complex social animals such as human and non-human primates. Lateral prefrontal cortex (LPFC) plays a key role in this ability, controlling a series of processes collectively defined as “executive functions”, which allow to select, organize, and optimize behaviors in order to reach specific intended goals [1–3]. Most classical studies on executive functions have examined the LPFC as a whole, without specific reference to its anatomical sectors [2,4–6]. However, the anatomical inhomogeneity [7–14] and functional complexity [15–21] of the LPFC suggest that it should not be viewed as a single anatomical and functional structure, but rather as comprising relatively distinct areas and sectors. For example, an anatomo-functional distinction between the dorsal and ventral sectors of the LPFC has been demonstrated by various studies [22,23]; in contrast, the presence of rostro-caudal anatomo-functional differences has been less investigated.

A series of fMRI studies in humans led to the hypothesis that executive functions are implemented by top-down control signals organized along a rostro-caudal axis in the frontal cortex. Koechlin and coworkers [15,24] proposed a ‘cascade’ model, in which the polar part of LPFC lies at the top of the hierarchy, being involved in maintaining relevant “pending” information about past events during multiple-task performance, the middle part of LPFC occupies an intermediate position, being involved in exploiting episodic information to be applied in a given behavioral context, while its posterior part uses the ‘immediate’ context-related information to select the correct action. Finally, in this model, the lateral premotor cortex, that lies on the other extreme of the hierarchy, is involved in coding stimulus-response associations. Recent studies confirmed the idea that progressively rostral LPFC areas are involved in processing more abstract information but attributed the “top” of this hierarchy to the middle part of LPFC rather than to its rostralmost part, suggesting that the middle LPFC is involved in linking both abstract and concrete (current) information for behavioral control [16,17,21,25].

The hypothesized rostro-caudal functional organization of the human LPFC is confirmed by data from patient studies showing that lesions of the lateral premotor cortex (BA6) were significantly associated with deficits in “sensory control”, lesions of the caudal prefrontal cortex (BA 45) with deficits in “contextual control”, while lesions of the rostralmost portion of LPFC (including BA 46 and 47) were associated with deficits in “episodic control” [26].

Connectional evidence in the monkey shows a rostro-caudal organization of LPFC, which can be subdivided into at least three vertical strips: a caudalmost strip, essentially connected with parietal and frontal areas and subcortical centers involved in oculomotor control [8,11,12,14,27–30]; an intermediate strip, connected with parietal and premotor areas and subcortical structures involved in grasping and reaching movements [8,11,12,14,27,30–32]; a rostral-most strip, showing mostly intrinsic prefrontal connections [8,11,14,27,30,31].

In line with this connectional inhomogeneity, recent electrophysiological studies described a similar functional rostro-caudal gradient. This was mainly observed in studies focused on working memory processes and tasks requiring saccadic movements as output responses [18–20,33,34]. In addition, this topographic organization can also be inferred from studies employing different type of motor behaviors, which showed that oculomotor responses are typically found in more caudal regions, while responses related to forelimb movements are usually found more rostrally within LPFC [35–39].

Although connectional studies and indirect electrophysiological evidence suggest the presence of distinct functional areas in LPFC, no electrophysiological study specifically focused on demonstrating the congruence between the distribution of specific functional properties and the cytoarchitectural and connectional parcellation. This may be due to the fact that most studies investigating the existence of distinct functional areas within the LPFC typically employed only a single task. The use of multiple tasks, probing the involvement of various LPFC areas in both passive processing of contextual information and its use to guide behavior, could be a more effective approach for identifying topographically organized functional differences, as suggested by the pioneering study of Tanila and coworkers [36].

To address these issues, in the present study we explored the potential functional inhomogeneity of LPFC, focusing on its ventral sector. To this aim, we recorded single neuron activity in monkeys performing three different tasks, investigating how visual information is either passively processed or exploited to guide a behavior involving the decision to produce or withhold grasping actions. Finally, we analyzed the data using the anatomical parcellations previously described by our group on the basis of cytoarchitectural and connectional studies [10,40].

## Results

The general aim of the present study was to identify the specific contribution of different prefrontal areas to visual processing and action organization. To this aim, 1) we recorded neuronal activity in the ventrolateral prefrontal (VLPF) cortex using three behavioral paradigms; 2) we identified distinct anatomical areas, based on architectural and connectional data; 3) we subdivided the neuronal responses on the basis of this anatomical parcellation and compared the functional properties observed in each area.

Concerning the behavioral paradigms, we employed two “passive” tasks aimed at assessing how VLPF neurons encode visual stimuli, in the absence of a specific request to use them, and a more complex Visuo-Motor task investigating functions related to action organization (see Methods; [38, 41, 42]). In particular, in the Picture task (Fig 1A), the monkeys simply had to observe one of 12 pictures; in the Video task (Fig 1B) they were required to observe one of 6 videos; in the Visuo-Motor task (Fig 1C), monkeys had to either grasp one of 3 objects (Action condition) or refrain from acting (Inaction condition), based on the color of instructing visual cues and the physical features of the presented objects.

**Fig 1.**
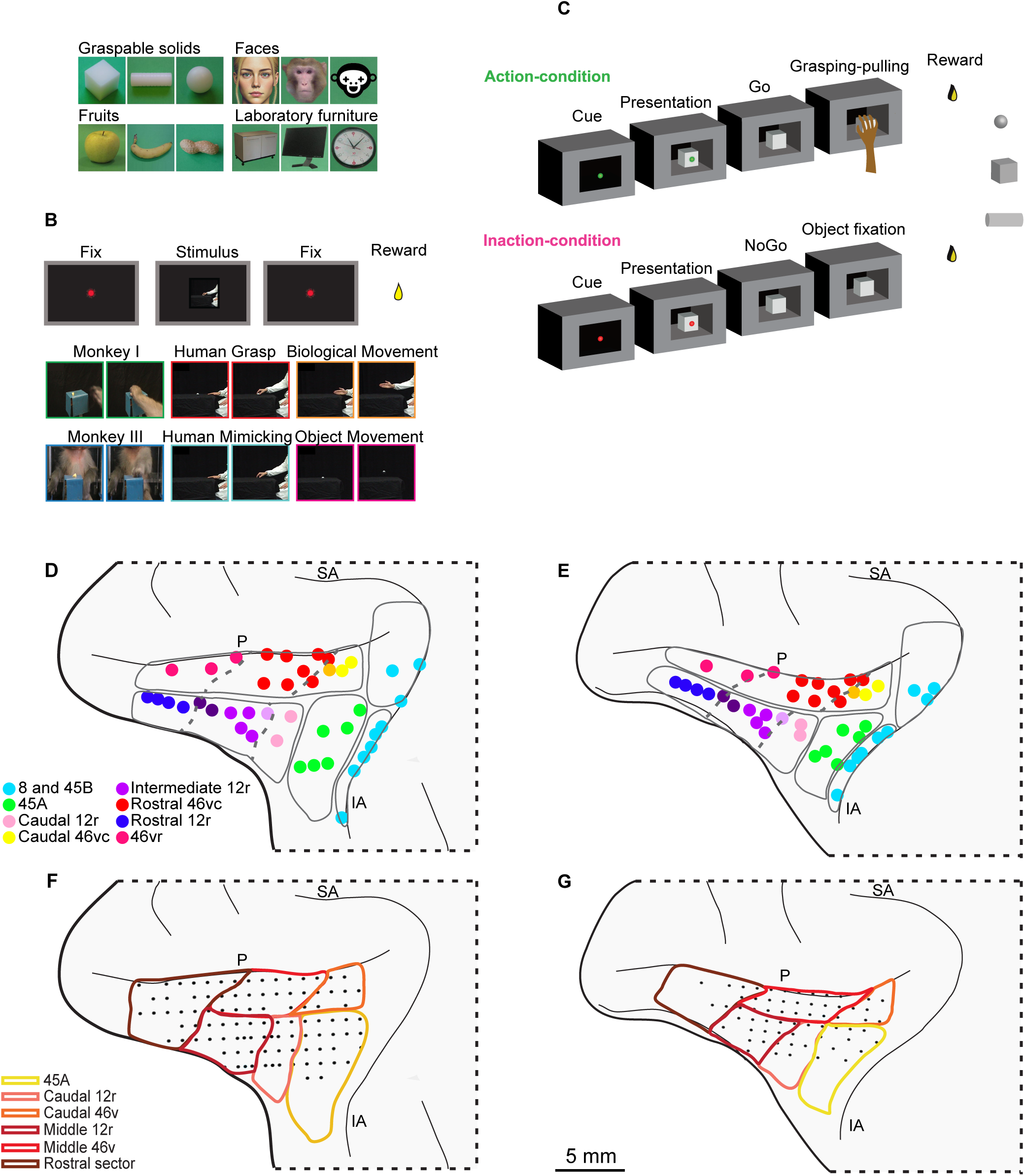
Behavioral paradigms and parcellation of the recorded regions. (**A** and **B**) Temporal sequence of events (above) and stimuli (below) of Picture and Video tasks. Note that the “human face” stimulus of the Picture task, depicted in this figure, has been here replaced by a DALL-E generated illustrative example for privacy reasons. **(C)** Temporal sequence of events of the Visuo-Motor task. On the right, the three used objects are shown. (**D** and **E**) Localization of the injection sites, taken from the literature (see Methods), superimposed on the cito-architectonically defined VLPF areas of the recorded monkeys. Each dot represents a different injection site. Each color refers to a specific pattern of connectivity characterizing the injection sites [11, 29, 31]. Note that, in the present work, we refer to caudal 46vc as caudal 46v, to rostral 46vc as middle 46v, to intermediate 12r as middle 12r and to 46vr as to rostral 46v (see Methods). Orange, light and dark violet dots refer to territories showing transitional connectivity. Dotted lines correspond to the borders between connectionally defined areas. (**F** and **G**) Recording grids of the two monkeys superimposed on the anatomical parcellation shown in (D) and (E). P: Principal sulcus; IA: Inferior arcuate sulcus; SA superior arcuate sulcus.

In order to define distinct VLPF areas in the recorded brains, we grouped the injection sites from previous connectional studies by our lab [43–45] based on their patterns of connection and warped their location on the histological reconstruction of the recorded brains (see Methods and *46)*. Figure 1 depicts the result of this process and the location and extent of anatomical areas of monkey 1 (M1, Fig 1D) and 2 (M2, Fig 1E). In this paper, we excluded from analysis the caudal oculomotor areas 8 and 45B and pooled together rostral 46v and 12r (hereafter defined as the *rostral sector*), since the position of the recording chambers allowed us to record only a small number of sites, especially in M2 (see Methods).

Within the investigated region, we recorded neural activity from 99 penetrations in M1 (Fig 1F) and 64 in M2 (Fig 1G). Note that, while the central part of the recorded region was densely and homogeneously explored in both monkeys, Area 45A has a wider grid in M2. Furthermore, in M2, the sampling of the rostral sector was limited to its caudalmost part.

### Distribution of VLPF responses recorded in the Picture task

We recorded a total of 2289 neurons in the Picture task (326 in 45A, 208 in Caudal 46v, 575 in middle 46v, 285 in caudal 12r, 558 in middle 12r and 337 in the rostral sector), and analyzed their responses by means of a 2X12 ANOVA (Factors: Epochs, Stimuli, p<0.01) followed by Newman-Keuls post hoc tests, which allowed us to identify neurons responding to the observation of static images and assess their possible selectivity (for the selection criteria, see Methods and *42*).

Neurons responding to the presentation of static images are well represented in the whole recorded region in both monkeys, except for area 45A, characterized by a lower number of responses in M1 than in M2 (Figs 2A, B).

**Fig 2.**
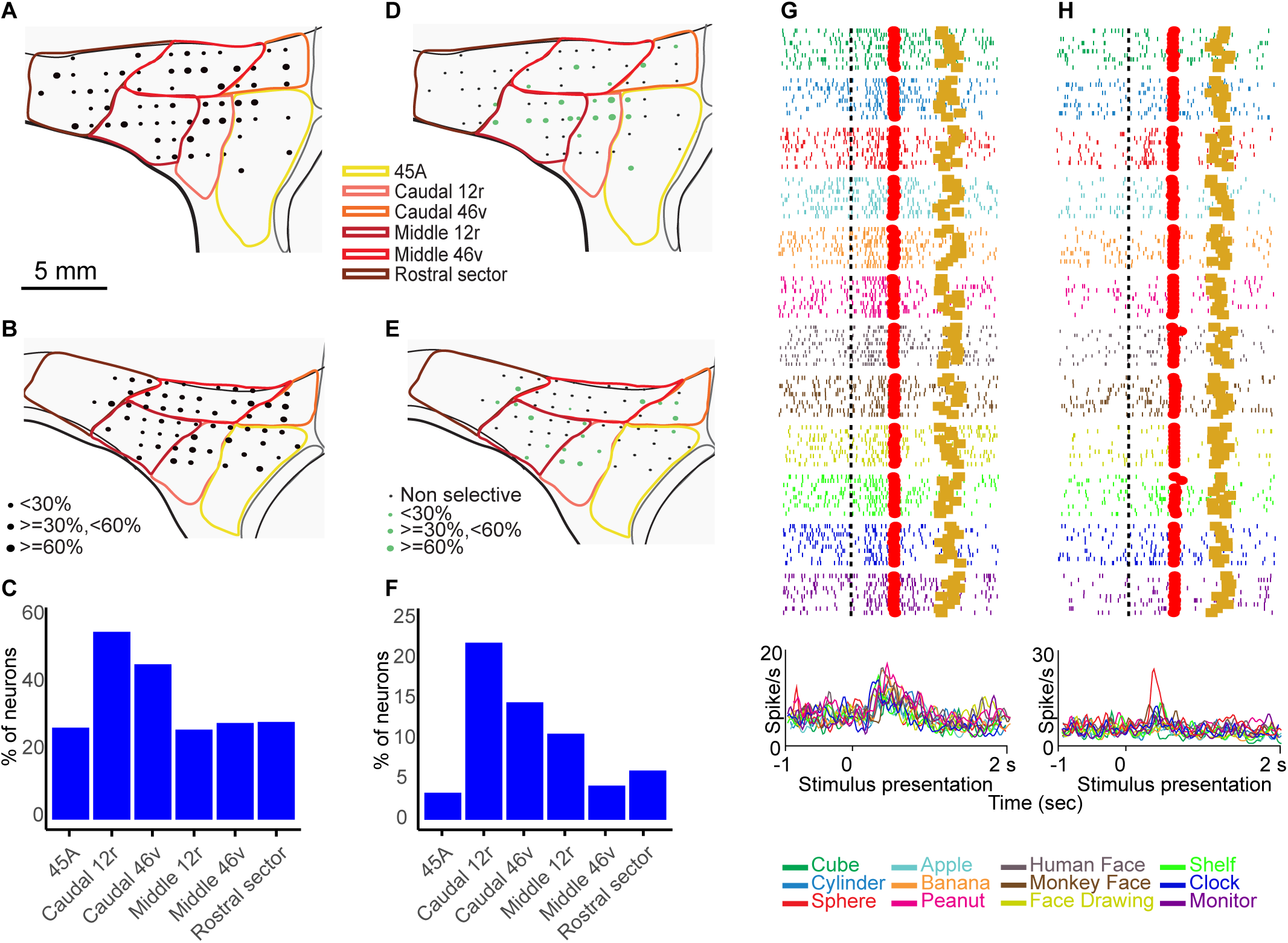
Distribution of functional properties in the Picture task. (**A**, **B, D, E**) Distribution of task-related neurons (above) and selective neurons (below) in the Picture task. The size of black and green dots represents, for each site, the proportion of task-related neurons on total neurons, or the proportion of selective neurons on task related neurons, respectively. (**C** and **F**) Barplots showing the percentage of task-related neurons on the total number of neurons (above) and of selective neurons on the total number of task-related neurons (below) recorded in each area, considering the two monkeys together. (**G**) Example of neuron showing a similar discharge profile following stimulus presentation in all conditions. (**H**) Example of neuron responding selectively to the presentation of the sphere. Rasters and histograms are aligned (vertical dashed line) with stimulus presentation. Rew squares: stimulus offset; Gold squares: reward delivery. Abscissae: time (s); Ordinates: firing rate (spikes/s).

Considering together the data of the two monkeys, caudal 46v and caudal 12r show the largest proportion of responsive neurons (Fig 2C). Selective neurons are mostly located in caudal 46v, caudal 12r and middle 12r (Fig 2D, E), being particularly represented in caudal 12r (Fig 2F). This distribution is also supported by the results of demixed principal component analysis (dPCA, see Methods). Indeed, during stimulus presentation, the stimulus identity explains the largest percentage of variance (see Methods) in these areas (caudal 12r: 27%; caudal 46V: 23%, middle 12r: 17%) than in the other considered areas (45A: 15%; rostral sector: 13%, middle 46v: 11%, S1 Fig). Figure 2G depicts the response of a caudal 46v non-selective neuron (2X12 ANOVA epoch effect p <0.01; interaction effect: n.s.). Fig 2H shows the response of a caudal 12r selective neuron, only active during the observation of the sphere (2X12 ANOVA interaction effect followed by Newman–Keuls post hoc test, p < 0.01).

Performing the same analysis on all recorded neurons we observed a very similar distribution. Indeed, the highest percentages were observed in caudal areas 12r and 46v and in middle 12r (26%, 20% and 16%, respectively), followed by the remaining areas (middle 46v:12%, 45A: 11%, rostral sector: 11%, S1 Fig).

### Distribution of VLPF responses recorded in the Video task

We recorded a total of 2687 neurons in the Video task (406 in 45A, 261 in Caudal 46v, 706 in middle 46v, 338 in caudal 12r, 632 in middle 12r and 342 in the rostral sector), and analyzed their responses by means of a 3X6 ANOVA (Factors: Epochs, Stimuli, p<0.01) followed by Newman–Keuls post hoc tests, which allowed us to identify neurons responding to the observation of the videos and assess their possible selectivity according to the criteria defined in the Methods section (see also [41]).

Figure 3A and B show the distribution of neurons responding to the observation of videos. In both monkeys, these neurons are more concentrated in the caudal and ventral part of the recorded region, being more homogeneously distributed in M1, and mostly located in caudal 46v and caudal 12r in M2. Neurons showing a preference for at least one stimulus have a similar pattern of distribution, being mostly located in the caudal and ventral areas (Fig 3D, E). Considering the two monkeys together, caudal areas 12r and 46v have the largest proportion of video responses (Fig 3C), with the latter being characterized by the highest proportion of selective neurons (Fig 3F). This distribution is in line with the results of dPCA (see Methods): considering the neuronal activity recorded during video presentation, the type of video explains the largest percentage of variance in caudal 46v (33%), followed by caudal 12r (27%). In the remaining areas, this factor explains a smaller percentage of variance (middle 12r: 22%; 45A: 19%; middle 46v: 16%; rostral sector: 17%, S2 Fig). Note that the same analysis performed on all recorded neurons revealed a very similar trend, for which the highest percentages were observed in caudal areas 46v and 12r (28% and 24%, respectively), followed by the remaining areas, showing relatively similar percentages (middle 12r: 16%, 45A: 15%, middle 46v:13%, rostral sector: 4%, S2 Fig).

**Fig 3.**
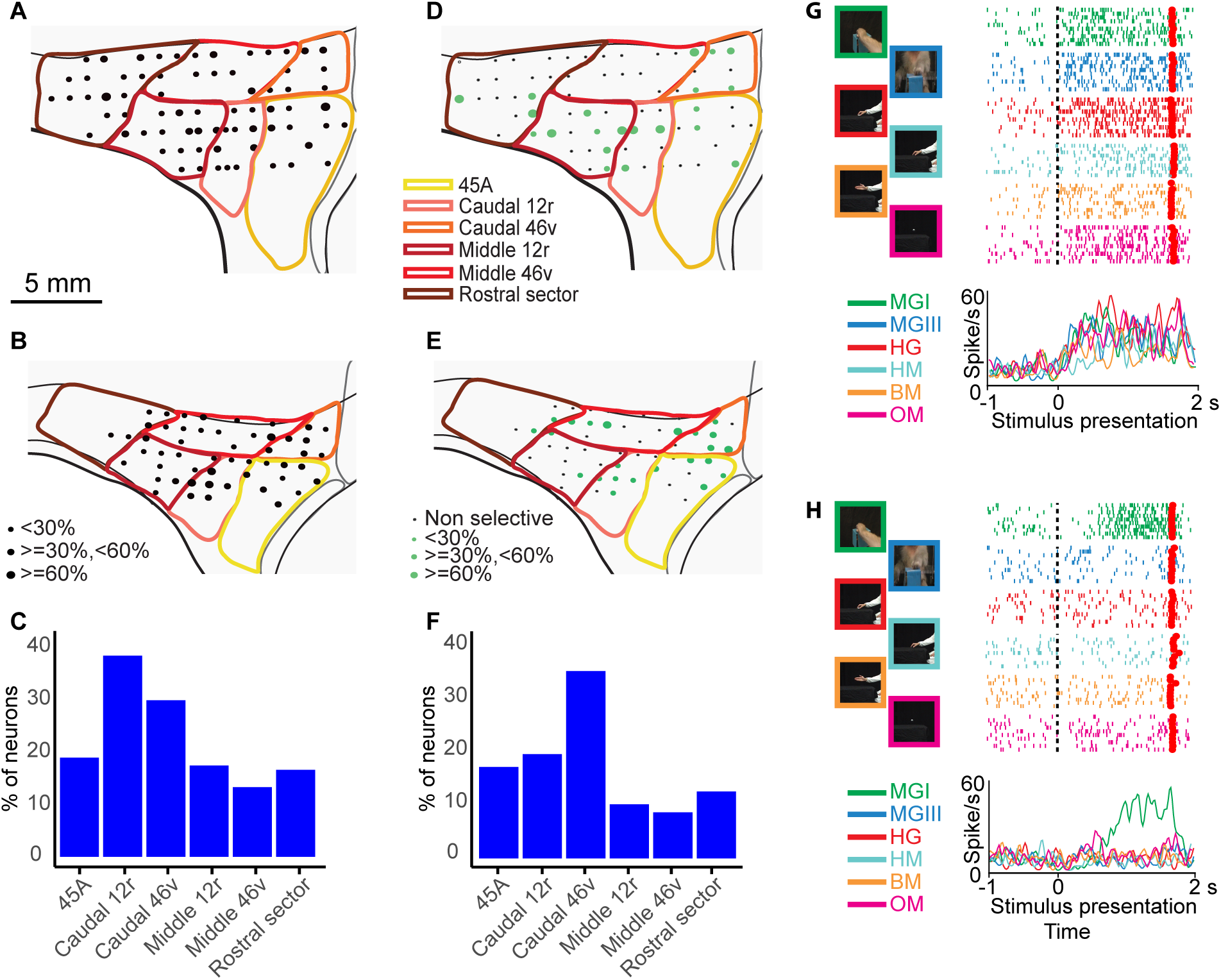
Distribution of functional properties in the Video task. (**A**, **B, D, E**) Distribution of task-related neurons (above) and selective neurons (below) in the Video task. (**C** and **F**) Barplots showing the percentage of task-related neurons on the total number of neurons (above) and of selective neurons on total task-related neurons (below) recorded in each area, considering the two monkeys together. (**G**) Example of neuron showing a similar discharge profile following stimulus presentation in all conditions. (**H**) Example of neuron responding selectively to presentation of a monkey grasping in first person perspective. Other conventions as in Fig 2.

Fig 3G shows an example of a caudal 12r non-selective neuron responding to video presentation, independent of the observed stimulus (3X6 ANOVA epoch effect, p < 0.01; interaction effect: n.s.). Fig 3H depicts the activity of a caudal 12r selective neuron discharging only during the observation of a monkey grasping food, seen from a third person perspective, and exclusively in the second video epoch (3X6 ANOVA interaction effect followed by Newman– Keuls post hoc test, p < 0.01).

### Distribution of VLPF single neurons responses recorded in the Visuo-Motor task

Using the Visuo-Motor task, we recorded a total of 3045 neurons (446 in 45A, 307 in caudal 46v, 761 in middle 46v, 392 in caudal 12r, 714 in middle 12r and 425 in the rostral sector). We analyzed the neuronal responses by means of a 9X2 ANOVA (Factors: Epoch, Condition, p<0.01), followed by Newman-Keuls post hoc tests, which allowed us to identify task-related neurons, and assess their possible preference for one of the two task conditions in each epoch, according to the criteria defined in the Methods section (see also [47]). Figure 4A and B depict the distribution of task-related neurons in the two monkeys. Note that task-related neurons are quite homogeneously represented in the different areas.

**Fig 4.**
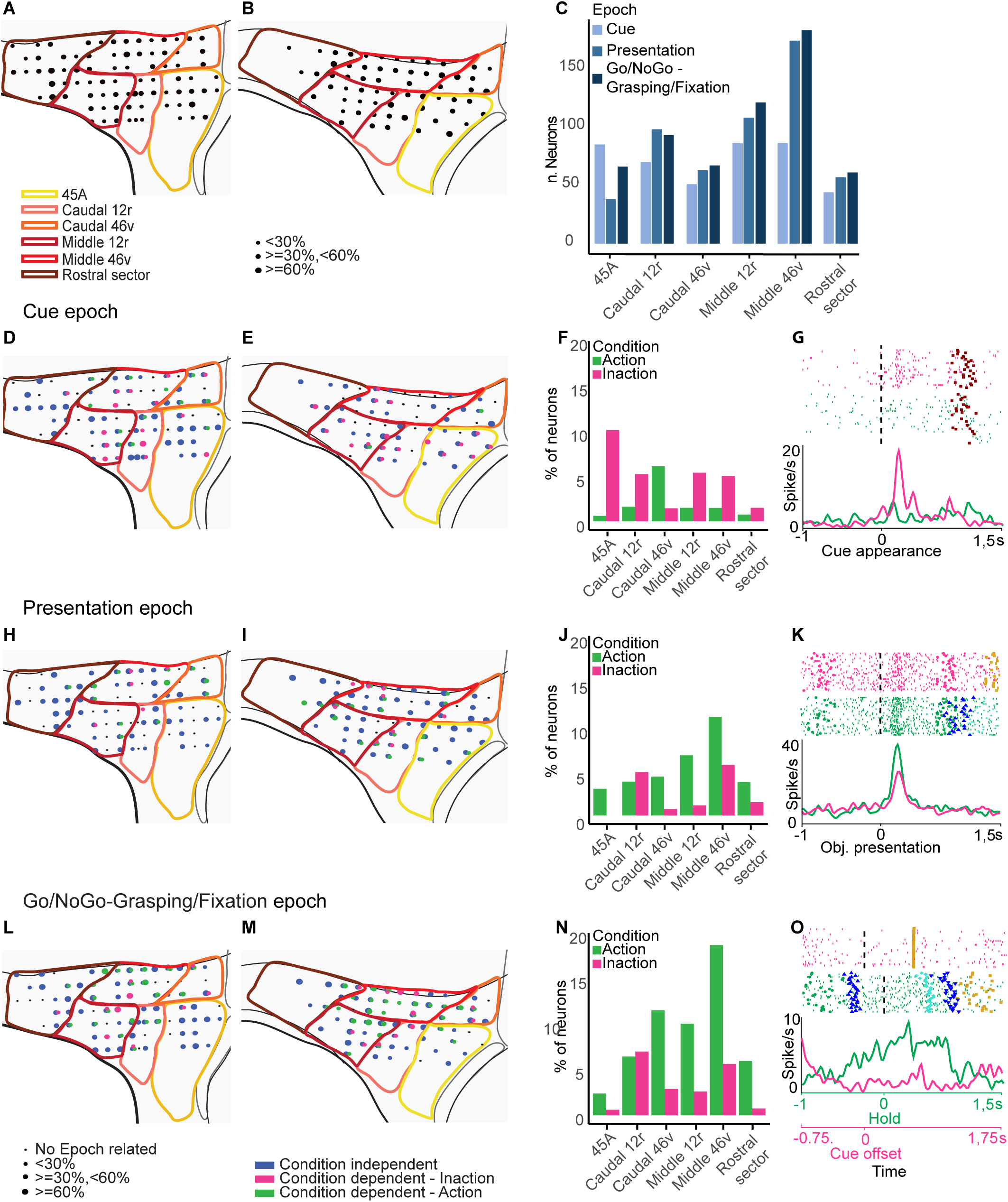
Distribution of functional properties neurons in the Visuo-Motor task. (**A** and **B**) Distribution of task-related neurons. (**D, E, H, I, L, M**) Distribution of Condition-dependent and Condition-Independent neurons responding during the Cue (second row) Presentation (third row) and Go/NoGo-Grasping/Fixation (fourth row) epochs. Black dots represent penetration sites in which no epoch-related neurons have been recorded. Blue, green and magenta dots correspond to the sites in which Condition-independent, Action selective or Inaction selective epoch-related neurons, respectively, are represented. The size of colored dots represents the proportion of the respective category of epoch-related neurons on the total number of task-related neurons observed in each site. (**F**, **J**, **N**) Barplots showing the percentage of Condition-dependent Cue-related (second row), Presentation-related (third row) and Go/NoGo-Grasping/Fixation-related (fourth row) neurons on the total number of task-related neurons recorded in each area, considering the two monkeys together. Green and magenta bars represent the Action and Inaction selective neurons, respectively. (**G**) Example of neuron showing a preferential discharge following cue appearance in the Inaction condition. (**K**) Example of neuron showing a discharge rate after object presentation, which is higher during the Action compared to the Inaction condition. (**O**) Example of neuron showing a discharge starting from the offset of the instructing cue, only during the Action condition. Rasters and histograms are aligned (vertical dashed line) with cue onset (**G)**, object presentation **(K**), and object holding/cue offset (**O**, for Action and Inaction conditions respectively). Green/Magenta circles: Action and Inaction cue appearance and offset, Brown squares: object presentation; Blue upward triangles: release of the hand from the starting position; Blue downward triangles: return of the hand on the starting position; Gold squares: reward delivery. Other conventions as in Fig 2.

Fig 4C shows the distribution of neuronal responses in the main task phases (see [47]) for each area, considering the two monkeys together. In 45A, the number of responses to cue appearance is higher than in the other periods. Caudal 46v and the rostral sector are characterized by a gradual increase in the number of responses from the beginning to the end of the task. A similar pattern, although associated with a higher number of responses in all considered periods, is present in middle 12r. Similarly, also caudal 12r and middle 46v are characterized by a generally high number of responses in all considered periods. Moreover, in these areas, the number of responses observed in the Presentation and Go/NoGo-Grasping/Fixation epochs is quite similar and much higher than that observed in the Cue epoch.

Based on the above-mentioned statistical analysis, we identified neurons responding, in each epoch, either preferentially to one of the two conditions (Condition-dependent, see Methods) or independently of the condition (Condition-independent).

*Cue-related* neurons are more densely represented in the ventral part of the recorded region than in the dorsal one in both monkeys, with neurons showing a preference for the Inaction condition being much more represented than those preferring the Action one (Fig 4D, E). Indeed, considering the two monkeys together, the proportion of neurons showing a preference for the Inaction condition is higher than that of neurons preferring the Action condition in all the considered areas, except for caudal 46v in which this trend is inverted (Fig 4F). The proportion of Inaction-related neurons is particularly high in areas 45A, caudal 12r, middle 12r and middle 46v. Figure 4G shows the discharge of a 45A Condition-dependent neuron responding to cue appearance only in the Inaction condition (9X2 ANOVA, interaction effect followed by Newman–Keuls post hoc test, p < 0.01). Finally, in the rostral sector, the proportion of Condition-dependent neurons is very low, while Condition-Independent responses are largely prevalent, especially in M1.

*Presentation-related* responses are well represented in each area, and their distribution is quite homogeneous in both monkeys. However, in M1 these responses are relatively less represented in the central part of the recorded region (corresponding to middle 46v and middle 12r) and in 45A (Fig 4H, I). Condition-dependent neurons are more densely represented in caudal 46v, middle 46v, caudal 12r and middle 12r than in 45A and the rostral sector. Considering the two monkeys together, the proportion of Action-related neurons is much higher than that of Inaction-related ones in all areas except caudal 12r, where the numbers are quite balanced (Fig 4J). Figure 2K shows the discharge of a middle 12r Condition-dependent neuron responding to object presentation in both conditions, but significantly stronger in the Action condition than in the Inaction one (9X2 ANOVA interaction effect followed by Newman–Keuls post hoc test, p < 0.01).

Neurons responding in correspondence of the last part of the task (*Go/NoGo-Grasping/Fixation* epoch) are well distributed along the whole explored region (Fig 4L, M), with a predominance of Condition-independent neurons in 45A and in the rostral sector, and of Condition-dependent neurons in the central part of VLPF (caudal 12r, caudal 46v, middle 12r, and middle 46v). Considering the two monkeys together, the proportion of neurons preferring the Action condition is very high especially in middle 46v followed by caudal 46v and middle 12r while caudal 12r appears to be characterized by a balanced representation of neurons showing a preference for the Action or Inaction conditions (Fig 4N). Figure 4O depicts the discharge of a middle 46v Condition-dependent neuron responding only in the Action condition (9X2 ANOVA interaction effect followed by Newman–Keuls post hoc test, p < 0.01). The neuron starts discharging just before the beginning of movement, peaks during object pulling, and abruptly ceases firing when the action is completed with the hand returning to the starting position.

### Population activity of VLPF areas in the Visuo-Motor task

In order to evaluate the response of the population of neurons recorded in the different VLPF areas, we plotted, for each of them, the mean-net activity (see Methods) of task-related neurons (Fig 5) and of all recorded neurons (S3 Fig) in the Action and Inaction conditions.

**Fig 5.**
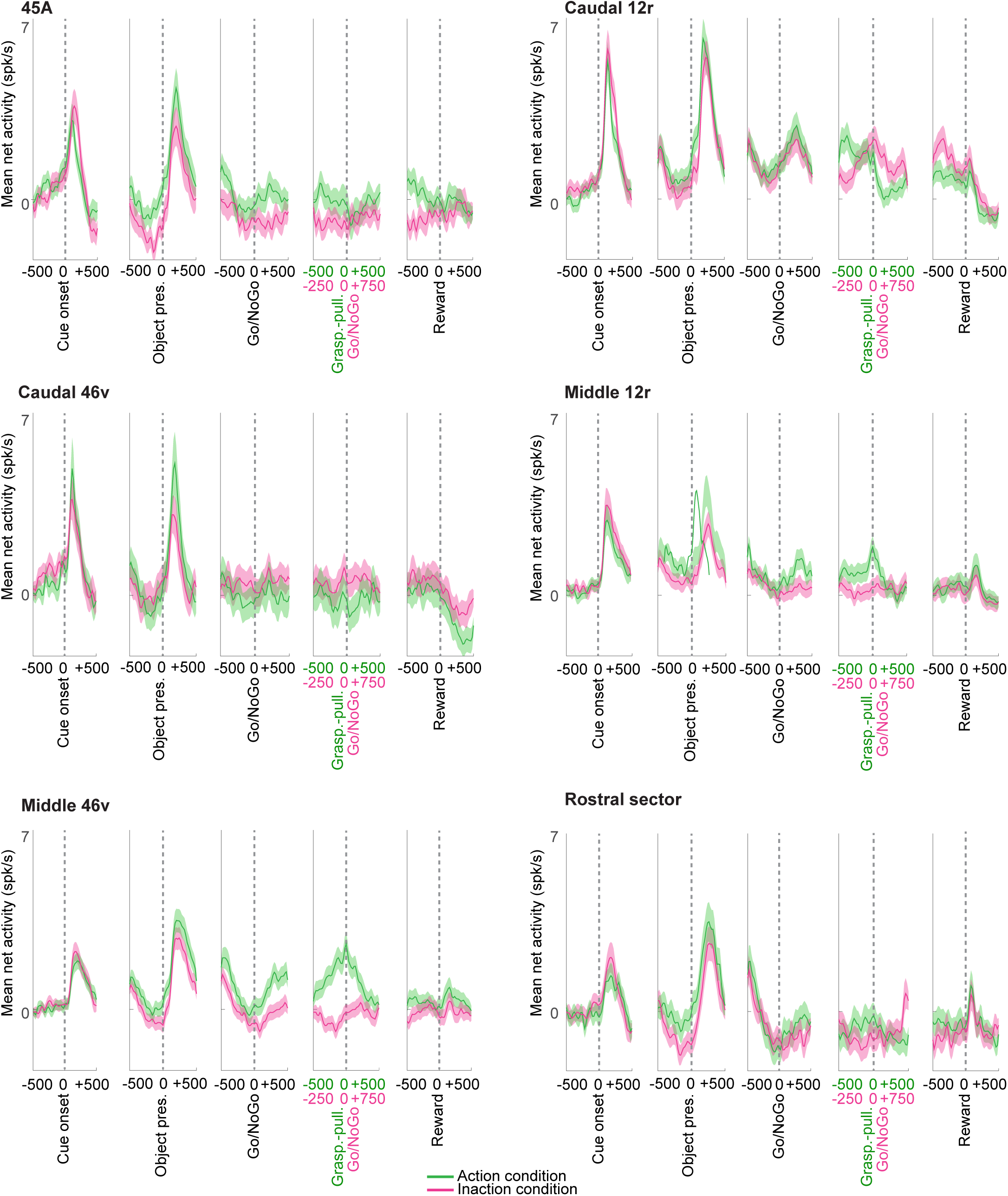
Mean population activity of neurons recorded in each area during the Visuo-Motor task. Temporal profile of mean net activity of the population of task-related neurons recorded in each of the considered VLPF areas. The magenta and green curves indicate the population mean net activity in the Inaction and Action conditions. The shaded area around each curve represents standard errors. The neuronal activity is aligned on the main task events indicated below each panel of the figure.

Both analyses showed that all areas share, as a common feature, the presence of two clear peaks following cue and object appearance. The peak related to cue presentation during the Inaction condition is higher than the one observed during the Action condition in all areas, except for caudal 46v, where this pattern is inverted. The object presentation-related peak is, in all areas, higher than the cue-related one, with the Action condition always eliciting a stronger population activity than the Inaction one during this period.

The pattern of responses observed in the final part of the task (starting around the Go/NoGo signal), is instead quite heterogeneous, when comparing the different areas (Fig 5 and S3 Fig). Some of them are mainly characterized by an enhanced population response, while others show a suppressed one (i.e. below baseline level). In particular, the rostral sector shows a suppressed population activity in both Action and Inaction conditions, while 45A shows activity suppression only in the Inaction condition, being the Action related response around baseline level. Caudal 46v exhibits a slight suppression of the activity occurring mainly in the Action condition just before the Go/NoGo signal. Note that, when considering the population of task-related neurons (Fig 5), the activity suppression observed in this condition is maintained also around object holding. Caudal 12r population activity starts growing in both conditions after the Go/NoGo signal, reaching a peak after about 250 ms. In the Inaction condition, the activity remains sustained until the reward delivery, then briskly falls below baseline level, while in the Action condition the sustained activity ends after the beginning of holding, abruptly decreasing to baseline level, and further decreasing below it, after reward delivery. Middle 12r is characterized by a sustained increase in firing rate only in the Action condition. The population response starts growing after the Go signal, reaching a peak around the beginning of holding. Then, the activity decreases to baseline level while the monkeys keep pulling the object. In both conditions, there is a final, lower peak after reward delivery. Middle 46v shows, in the Action condition, the highest mean-net activity among all the considered areas in the final part of the task. Notably, while in the Inaction condition no clear enhancement or suppression is evident, in the Action condition the activity grows after the Go signal, continues to increase during reaching and grasping execution, peaks around the beginning of holding, and gradually decreases while the monkeys hold the object, returning to baseline level around reward delivery.

### Condition and Object dependency in the VLPF areas

In order to evaluate how neurons of each area encode the Condition and Object factors during the unfolding of the Visuo-Motor task, we adopted a data-simplification method: the demixed principal component analysis (dPCA) on specific time periods (Cue, Presentation and Go/NoGo-Behavioral response, see Methods).

Concerning the Cue period, we used only the Condition factor, since the Object one is still not present in this phase of the task. Performing the analysis on task-related neurons, we observed that caudal 46v shows the highest percentage of variance explained by the Condition factor (27 %), followed by middle 46v (23%), caudal 12r (21%), middle12r (20%,) and 45A (14%), while the rostral sector shows the lowest percentage of condition-dependent variability (13%). Concerning the Presentation period, we used both Condition and Object factors for the analysis. Considering task-related neurons, in all the considered areas apart from caudal 12r and the rostral sector, the percentage of variance explained by the Condition factor is higher than that explained by the Object factor. Similarly to Cue epoch, the percent of signal variance explained by the Condition is the highest in middle 46v (22%) followed by caudal 46v (17%), caudal and middle 12r (13% and 11%), rostral sector (9%) and 45A (7%). In all areas, except caudal 12r and 45A, the Condition effect is already significant at the object appearance. The percentage of variance explained by the Object factor is the highest in caudal 12r (15%) followed by caudal 46v and rostral sector (11%, 10%), middle 12r and middle 46v (both 8%) and the lowest in 45A (4%) (S5 Fig). In all areas (except 45A, which shows no significant effect) we observed a significant Object effect emerging with a timing comparable with the development of the population response to object appearance (compare S5 and S7 Figs).

Regarding the Go/NoGo-Behavioral response period, when analyzing task-related neurons we found that in caudal 12r the percentage of variance explained by the Condition and Object factors are quite balanced (11% and 13%), while in the remaining areas the Condition factor is more represented than the Object factor (see S6 Fig). The highest percentage of variance explained by the Condition factor is reached by middle 46v (29%) followed by 45A (18%), caudal 46v (16%), middle 12r (14%), and rostral sector and caudal 12r (11%). Note that in middle 46v, caudal 12r and middle 12r, the Condition effect is already significant at the beginning of the period, while in the remaining areas it develops later. The percentage of variance explained by the Object factor is the highest in caudal 12r and caudal 46v (34%) followed by middle 12r (27%), rostral sector (25%), 45A (13%) and middle 46v (9%) (S6 Fig). All the observations described above on the populations of task-related neurons analyzed for each area, were generally confirmed when performing dPCA on all neurons recorded in each area (S7-S9 Figs).

### Decoding of the condition and object factors in VLPF areas

We adopted a decoding approach to assess, in a time-detailed manner, which type of information is encoded in the distinct time epochs of the Visuo-Motor task by the studied VLPF areas. Figure 6 shows the accuracy level of the decoding of the Condition (A-F) and the Object (A’-F’) factors when the classifier is trained on a specific time period and tested on all time periods (cross-temporal decoding plots, see Methods).

**Fig 6.**
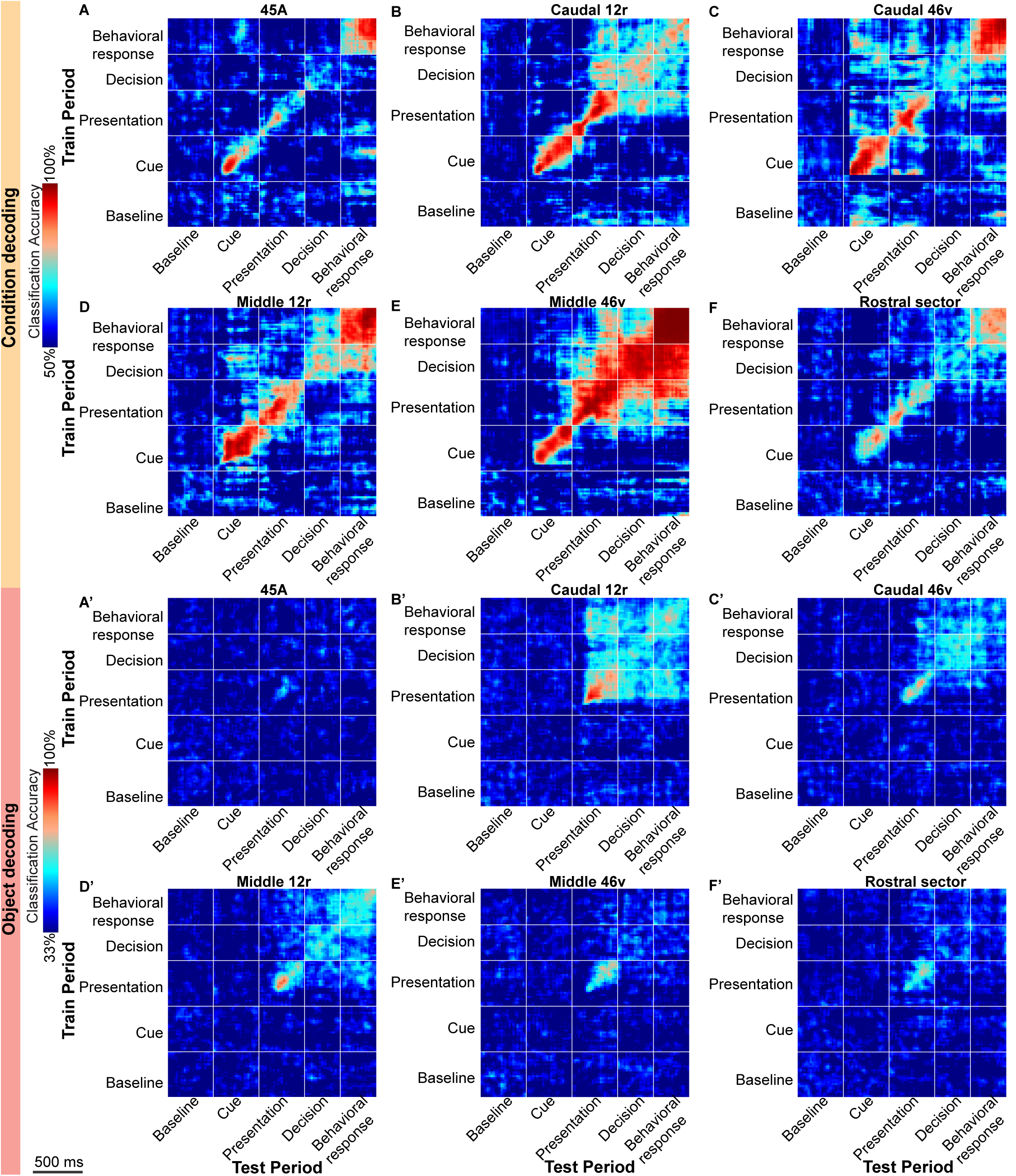
Cross-temporal decoding of the Condition (A-F) and Object (A’-F’) factors of the Visuo-Motor task in task-related neurons. For each analysis, the decoding accuracy is computed in bins of 60 ms, sampled at 20 ms intervals. For each plot, the vertical and horizontal lines delimit the considered time periods (see Methods). Decoding periods of testing and training are indicated on the X and Y axes, respectively.

As in the previous cases, the decoding analyses were performed considering both the population of task-related neurons and that of all recorded neurons within each area. Since the results were strongly consistent across these analyses, they will be described together. For all areas, considering the Condition factor, the highest accuracy is reached when the classifier is trained and tested on the same time bins, i.e. on the diagonal, and typically the lowest level along the diagonal occurs in the Decision period. Taking into account the Object factor, the highest accuracy is always observed, in each area, along the diagonal in the Presentation period. In the following paragraphs we will focus on the features characterizing each area.

#### 45A

In area 45A, the accuracy observed along the diagonal reaches three distinct peaks in the Cue, Presentation and Behavioral response periods. The decoding performance is also high when training on data from the final phase of the Cue period and testing on data from the final part of the Behavioral response and vice versa, indicating that there is a static pattern of activity (see Methods) representing each condition across these two time periods (see Fig 6A for the result on task related neurons and S12A Fig for the results on all recorded neurons). Note that this pattern is statistically significant (cluster-based permutation test, p<0.001, see S10A and S13A Figs and Methods). Another significant static pattern of activity is observed when training the algorithm on the very early part of the Presentation period and testing on the Behavioral response period (Fig 6A, S10A, S12A and S13A Figs).

The decoding of the Object factor reveals a quite low, though significant, accuracy level restricted to the Presentation period (permutation test p<9.5e-6, Bonferroni corrected for the number of on-diagonal time bins Fig 6A’, S11A, S12A’ and S14A Figs).

#### Caudal 12r

Considering caudal 12r, the decoding of the Condition factor reveals that, along the diagonal, the accuracy levels are higher during the Cue and Presentation periods and lower (although still around 75% and significant) in the final phases of the task (permutation test p<9.5e-6, Bonferroni corrected for the number of on-diagonal time bins; Fig 6B, S10B, S12B and S13B Figs). Considering the cross-temporal decoding, each condition is represented by a pattern of activity which is shared among the Presentation, Decision and Behavioral response periods (static pattern; cluster-based permutation test, p<0.001; Fig 6B, S10B, S12B and S13B Figs).

The decoding of the Object factor reveals, along the diagonal, a statistically significant accuracy from the Presentation period until the end of the task (permutation test p<9.5e-6, Bonferroni corrected for the number of on-diagonal time bins; Fig 6B’, S11B, S12B’ and S14B Figs). The accuracy is the highest in the Presentation period, decreases in the Decision and slightly increases again in the Behavioral response period (Fig 6B’). Cross-temporal decoding shows that the different objects are represented by specific patterns of activity, shared across the Presentation, Decision and Behavioral response periods (static pattern, cluster-based permutation test, p<0.001, Fig 6B’, S11B, S12B’ and S14B Figs).

#### Caudal 46v

Caudal 46v is characterized by a very high and significant Condition decoding accuracy occurring, along the diagonal, in the Cue, Presentation and Behavioral response periods, while the Decision period is characterized by lower, but still significant, accuracy levels (permutation test p<9.5e-6, Bonferroni corrected for the number of on-diagonal time bins; Fig 6C, S10C, S12C and S13C Figs). Concerning cross-temporal decoding, a significant static pattern of coding is observed in several phases of the task (cluster-based permutation test, p<0.001; Fig 6C, S10C, S12C and S13C Figs). In particular, the two conditions are represented by patterns of activity, which are shared among 3 time periods: Cue, Presentation and a phase extending from the end of the Decision period to the beginning of Behavioral response one (static pattern; cluster-based permutation test, p<0.001; Fig 6C, S10C, S12C and S13C Figs). Note that this result is more evident when training the classifier on the last part of the task and testing it on the Cue and Presentation periods, and less evident in the other direction.

Decoding of the Object factor reveals, along the diagonal, a statistically significant accuracy from the Presentation period until the end of the task (permutation test p<9.5e-6, Bonferroni corrected for the number of on-diagonal time bins; Fig 6C’, S11C, S12C’ and S14C Figs). Cross temporal decoding shows a similar static pattern as that described for area caudal 12r (cluster-based permutation test, p<0.001; Fig 6C’, S11C, S12C’ and S14C Figs), although with slightly weaker accuracy levels.

#### Middle 12r

Condition decoding reveals, in middle 12r, a continuous and very high level of accuracy occurring, along the diagonal, during the Cue and Presentation periods, which decreases in the Decision period and raises again at very high levels in the Behavioral response period, remaining significant throughout all this period (permutation test p<9.5e-6, Bonferroni corrected for the number of on-diagonal time bins; Fig 6D, S10D, S12D and S13D Figs). Cross-temporal decoding reveals that the two conditions are represented by patterns of activity that are shared across the Cue, Decision and Behavioral response period (cluster-based permutation test, p<0.001; Fig 6D, S10D, S12D and S13D Figs), especially when training the algorithm on the Decision period.

The decoding of the Object factor shows results similar to those described for caudal 12r and caudal 46v (static pattern, cluster-based permutation test, p<0.001, Fig 6D’, S11D, S12D’ and S14D Figs), although with slightly weaker accuracy levels considering the cross-temporal decoding results (cluster-based permutation test, p<0.001; Fig 6D’, S11D, S12D’ and S14D Figs).

#### Middle 46v

The decoding of the Condition factor in middle 46v reveals, along the diagonal, a continuous, significant and very high level of accuracy starting from the early phase of the Cue period until the end of the task (permutation test p<9.5e-6, Bonferroni corrected for the number of on-diagonal time bins; Fig 6E, S10E, S12E and S13E Figs). Cross-temporal decoding results reveal a common pattern of activity representing the Condition factor during most of the task, including Presentation, Decision and Behavioral response periods (cluster-based permutation test, p<0.001; Fig 6E, S10E, S12E and S13E Figs).

The decoding of the Object factor reveals that a significant and quite high level of accuracy is observed almost exclusively along the diagonal, in the Presentation period (permutation test, p<9.5e-6, Bonferroni corrected for the number of on-diagonal time bins; Fig 6E’, S11E, S12E’ and S14E Figs).

#### Rostral sector

Although the decoding of the Condition factor in the rostral sector shows the lowest levels of accuracy among the studied areas, relatively high levels of accuracy are reached, along the diagonal, in the Cue, Presentation and Behavioral response periods, while the Decision period is characterized by a lower but still significant level of accuracy (permutation test p<9.5e-6, Bonferroni corrected for the number of on-diagonal time bins Fig 6F, S10F, S12F and S13F Figs). A significant static pattern of Condition coding, though associated to a quite low level of accuracy, is observed only when training the classifier on the Decision period and testing it on the Behavioral response one, and vice versa (cluster-based permutation test, p<0.001; Fig 6F, S10F, S12F and S13F Figs).

The decoding of the Object factor reveals that a significant and quite high level of accuracy is observed exclusively along the diagonal, in the Presentation period (permutation test, p<9.5e-6, Bonferroni corrected for the number of on-diagonal time bins; Fig 6F’, S11F, S12F’ and S14F Figs).

## Discussion

In the present study, we investigated the specific contribution of different monkey VLPF areas in processing sensory information and exploiting it for guiding behavior. To this aim, we recorded neuronal activity from a large sector of VLPF, covering about its posterior two thirds, during tasks involving either in the passive processing of visual stimuli or their use for guiding reaching-grasping actions.

Under the hypothesis that the connectional features of different areas underpin a functional specificity, we attributed the location of recorded neurons to different connectionally defined anatomical areas, showing that the distribution of functional properties largely matches the adopted hodological parcellation.

Our main results show that:

- The passive presentation of visual stimuli primarily activates neurons in the caudal VLPF areas, particularly in the caudal 12r. This suggests that these regions represent the first VLPF processing stage of visual input, consistent with their strong connections to the inferotemporal cortex [31];
- The execution or withholding of grasping actions predominantly activates neurons in the intermediate VLPF areas, particularly in middle 46v. This indicates that these regions may primarily contribute to the selection and guidance of contextually appropriate actions, in line with their strong connections to the parietal and premotor cortices [11];
- Demixed principal component analysis (dPCA) of neural activity reveals that both the task rule (behavioral condition) and the type of stimulus modulate activity across all recorded areas, suggesting that the entire investigated region is involved in executive functions. Furthermore, dPCA indicates that each subdivision is characterized by distinct functional features.
- Decoding analyses on the neural activity recorded in the visuomotor task show that each VLPF area has a shared neural code across specific epochs of the task, suggesting that each area plays a distinct role in encoding the contextual information relevant for behavior.

### Comparison with previous mapping studies

In the past, several studies tried to functionally subdivide the lateral prefrontal cortex (including VLPF and DLPF) by using either passively presented visual stimuli or the production of movements in a context of working memory [18,20,33,48]; however, these tasks were usually examined in isolation from one another. An exception is represented by the work of Tanila and colleagues, who studied neural activity in a large peri-principal region during the presentation of visual, auditory, and somatosensory stimuli, as well as during the execution of hand and eye movements in a naturalistic context [36]. Interestingly, they found that visual responses were most densely represented in caudal VLPF, where also oculomotor neurons have been recorded, and somatic responses (i.e. those to somatosensory stimulation and/or during the execution of reaching-grasping actions) were primarily found in the intermediate VLPF. Accordingly, they suggested the presence of a functional distinction among different prefrontal sectors, based on their connectional properties. On the one hand, our results largely confirm their pioneering data. On the other, they provide a more refined and detailed description of how the functional properties of VLPF neurons are differentially distributed. Indeed, Tanila and collaborators adopted an anatomical parcellation based on large injections encompassing more than one area [49–54] and matched only a posteriori their functional data with this anatomical framework. In our study we instead relied on a more recent and detailed anatomical parcellation and used it for guiding our subsequent analyses. Finally, on the functional side, we used strictly controlled rather than more naturalistic paradigms.

The topographic distribution of visual responses in LPFC has been described more recently in a series of studies performed by the group of Constantinidis [18,19,34]. Although it is difficult to directly compare our data with their findings, mainly due to the use of markedly different anatomical frameworks, some clear similarity can be found. Indeed, in a recent study [19], the authors compared the responses to passive presentation of visual stimuli with those elicited by the same stimuli when used for a working memory task. They demonstrated that: posterior regions are typically more involved than anterior regions in coding visual stimuli during stimulus presentation and delay phases of the tasks, independently of whether the task was passive or memory related, while anterior areas are typically more involved in the memory task than in the passive [19]. In other words, these authors described the presence of a rostro-caudal functional gradient within the prefrontal cortex, in which posterior areas are more involved in coding visual information per se, while more anterior regions have a stronger role in exploiting visual information to guide behavior. Our results confirmed this type of functional organization and allowed us to extend this idea from working memory to action organization.

### The posterior region of VLPF constitutes an access node for visual information

The data from our visual tasks show that caudal and ventral VLPF areas are strongly involved in processing passively presented visual stimuli, independently of any association with a behavioral outcome. More specifically, our results show that caudal 12r, caudal 46v and, to a lesser extent, middle 12r are characterized by a high percentage of neurons showing a response during stimulus presentation and, as shown by dPCA, a high percentage of variance explained by the type of presented stimulus. In addition, these areas are characterized by the highest percentages of selective neurons. These functional properties strongly resemble those described by previous electrophysiological studies on caudal prefrontal areas [19,34,48,55] and high order temporal visual areas [56,57] and are in line with the fact that caudal 12r, caudal 46v and middle 12r show strong connections with temporal visual areas [10,11,31]. Thus, these prefrontal areas represent the first stage of processing of the visual input reaching VLPF.

Among caudal areas, 45A appears less involved in visual processing. Indeed, in this area we recorded a smaller number of responses to passively observed visual stimuli than in caudal 12r and caudal 46v. Noteworthy, this area has strong connections with posterior temporal areas providing visual and auditory information, including communicative stimuli, and with prefrontal regions and subcortical structures involved in oculomotor control and gaze orientation[28,29,58]. Thus, it is possible that the lower number of visual responses in this area with respect to the other caudal VLPF regions is due to the fact that neurons in this region typically respond to acoustic stimuli or to visual-acoustic stimuli combination [59,60]; these stimuli were not used in our paradigm.

### The intermediate region of VLPF exploits visual information for guiding actions

The rationale of our Visuo-Motor task allowed us to observe that the middle part of the investigated region is deeply involved in exploiting visual information for action guidance. More specifically, middle areas 46v and 12r show the largest number of neurons responding to object presentation, most of which are tuned to the Action condition. This evidence is in line with the functional gradients observed in literature [19,34], showing that these prefrontal regions have a stronger involvement in coding visual stimuli when these latter are actively exploited in relation to a subsequent behavioral response. Noteworthy, these areas are also characterized by the highest percentage of neurons specifically responding in the final phases of the task, during action programming and execution, in line with their strong connections with the parieto-premotor nodes of the grasping circuit [11,31,61].

### VLPF areas show overlapping yet partially different contributions to action organization

The variety of analysis we used to study the responses of neural populations allowed us to confirm the presence of a rostro-caudal functional gradient throughout the VLPF.

In particular, decoding analyses showed that caudal areas 46v and 12r and, to a lesser extent, middle 12r present a clear pattern of object coding which is shared between the phases of the task going from object presentation onward, indicating that caudal and ventral VLPF areas are more strongly involved in coding object-related visual information.

Instead, middle 46v and, to a lesser extent, middle 12r and caudal 46v, are characterized by strong condition-related effects (dPCA) and, as observed with the decoding analysis, by a very high accuracy in discriminating between conditions, with static patterns of activity representing the two conditions in the initial and final phases of the task (see below). This indicates that the intermediate/dorsal areas of VLPF are strongly involved in encoding the behavior associated to the actual context.

Finally, the rostral sector does not appear to participate to the functions investigated by our tasks (see also below).

Within this general gradient, we were also able to observe some peculiarities characterizing each investigated area.

Concerning Caudal 46v, decoding analysis shows that this area is characterized by a pattern of Condition coding that is shared between the Cue and Behavioral response periods. A similar pattern has been described in our previous work [47] for the population of neurons recorded in the whole VLPF preferentially responding to the cue appearance in the Action condition. We propose that this pattern indicates that the presented cue is already coded in terms of behavioral outputs (‘pragmatic’ hypothesis, see [47]). Looking at the mean population response, neurons of Caudal 46v show a preference for the Action condition during cue appearance and a small inhibition, in the same condition, around the Go/NoGo signal. Considering only task-related neurons, activity suppression is observed, in the same condition, also around object holding, when the eyes are actually free to move. Considering that this area is strongly connected with oculomotor centers [11,28], this inhibition suggests that the observed prospective coding could be linked to the oculomotor control, that is especially relevant in the final phases of the Action condition.

Considering middle 46v, the decoding analysis shows a very stable representation of the Condition from the Presentation period onwards, and dPCA shows that this area has the highest percentage of variance explained by the Condition, especially in the final phase of the task. In addition, the mean population response observed in this area revealed a clear preference for the Action condition from object presentation to the end of the task. The same pattern of activity has been described in our previous work [47], when considering the population of neurons recorded in the whole VLPF showing a preference for the Action condition. Altogether, these data indicate that neurons of middle 46v encode information related to the presented visual stimuli in terms of the subsequent goal to be achieved rather than in visual terms [2, 62–64]. In other words, neurons of middle 46v encode instructions and objects in pragmatic terms, namely as predictions of the critical events signaling the behavioral outcome (e.g., for example, taking possession and pulling the object and reward delivery, ‘pragmatic’ hypothesis; [47]).

These results are in line with electrophysiological studies showing that prefrontal neurons encoding the behavioral output during the execution of a motor task are located mainly in the intermediate portion of area 46v [36–38,65], as well as with its strong parietal and premotor connectivity [11,14,27]. These connections could provide middle 46v with the motor signals and sensory (somatosensory and visual) information necessary for the on-line control of behavior and for monitoring the achievement of the final action goal.

Concerning middle 12r, Condition decoding revealed a shared representation of this factor between Cue and Decision periods. A similar pattern was identified by Rozzi and coworkers [47] when analyzing the population of Cue-related neurons showing a preference for the Inaction condition. In line with the “pragmatic coding hypothesis”, this implies that neurons of middle 12r encode the cue appearance in the Inaction condition in terms of the expected feedback (that is keeping the hand on the starting position) signaling goal achievement.

When performing Object decoding, caudal and middle 12r show a similar type of static pattern, starting from Object presentation. This similarity is corroborated by dPCA results, showing that both areas are characterized by an early and strong encoding of presented objects. These two areas are connected with inferotemporal regions involved in the pictorial description of objects, as well as with parietal area AIP, which plays key roles in pragmatic coding of object features [31,66,67]. In addition to these connections, they are also connected with premotor areas involved in executing hand-arm actions and with caudal VLPF oculomotor areas responsible for generating eye movements [31]. Both 12r sectors are also connected to middle 46v, which, as previously noted, may contribute to on-line behavior control.

Altogether, the data on middle and caudal 12r suggest that, although both areas are involved in processing object related information, caudal 12r appears to mainly encode objects in visual terms, while middle 12r appears to play a major role in exploiting object information for guiding behavior. Area 45A is the only sector in which Cue responses are more represented than those to the other periods of the task. In addition, the mean population response is strongly enhanced during cue and object presentation and slightly inhibited during fixation maintenance (Behavioral response period of the Inaction condition). The nonspecific visual responses of neurons of this area to behaviorally relevant information could be related to possible attentional processes, in line with a series of observation indicating a specific involvement of this region in attention-related functions linked to the visual search of specific targets [68,69].

The rostral sector of the investigated region of VLPF is much less responsive to the tasks employed in this work. This does not appear to depend on the low number of recorded neurons, since their number is quite similar to that observed in the other investigated areas. The low responsiveness could depend on the fact that our tasks were not suitable to investigate the functions of this region. This negative result, however, allowed us to define a clear border between the above-described areas and this rostral territory.

The time course of the population response of this area, is characterized, in the Visuo-Motor task, by an interesting functional feature: the activity is suppressed during the whole task unfolding, in both Action and Inaction conditions, except for two excitatory peaks in correspondence of sensory stimuli presentation. This inhibition could be interpreted in light of a large corpus of literature showing that the rostral prefrontal cortex is deeply involved in several operations, such as coding behavioral rules based on past episodic information, coding self-generated decisions and storing conscious action plans [15–17,70,71]. As the task strongly depends on a clearly defined and immediately available context and is well learned by monkeys, the VLPF areas active during both Action and Inaction conditions could in the meantime inhibit the activity of rostral VLPF. This explanation is corroborated by the known strong intra-prefrontal connectivity of the rostral VLPF sector with more posterior VLPF regions [31,72].

### Limitations and further developments

Usually, neurophysiological studies test functions of cortical regions by using a task specifically aimed to evaluate the effect of few variables and by limiting the analysis to task-related neurons. In this paper, we employed three different tasks to better evaluate multiple functional variables on the same neurons. Furthermore, although in the main text we present the properties of task-related neurons, we also demonstrate that the results are confirmed, for each area, in the whole population of recorded neurons (see Supporting information). Thus, we can infer that the properties of the population of task-related neurons are representative of those of the whole investigated region within the studied functions.

However, the present work has, as a main limitation, the fact that the employed tasks only partially allowed us to characterize the functions of 45A and of the rostral sector of VLPF. In the present work, we tried to perform a functional parcellation of VLPF, by defining the role of different areas in context-based behavioral selection. Further studies could allow to better define the properties of these two cortical sectors, by verifying their response to other types of sensory stimuli (e.g. acoustic, somatosensory or multimodal), by assessing their role in high level guidance of oculomotor behavior, and in the processing of abstract information stored in episodic memory.

The types of motor behavior studied in this work, i.e. grasping objects and withholding actions, are daily performed also by humans, therefore our results could be important for verifying whether a modular organization of cognitive-motor functions similar to that here described in the monkey VLPF, is present also in our species. Interestingly, in many prefrontal neuropsychological syndromes, such as utilization behavior, echopraxia, and anarchic hand syndrome, the impairment of cognitive aspects is intrinsically linked to the motor ones [73–75]. Thus, we believe that the detailed anatomical identification of the prefrontal areas involved in different aspects of executive functions, such as context-based organization of behavior, will allow a more precise “mapping” of the symptoms of prefrontal syndromes.

## Materials and Methods

### Experimental Design

The general aim of the present study was to identify the specific role of different prefrontal areas in coding visual stimuli and exploiting this information for behavioral guidance. To this aim we recorded neuronal activity in two passive observation tasks and one Visuo-Motor task, and compared the neuronal responses VLPF areas, identified on the basis of architectural and connectional parcellations.

The experiment was carried out on two female Rhesus monkeys (*Macaca mulatta*, M1, M2) weighing about 4 kg. The animals have been previously employed in a series of experiments, whose results have already been published [38,41,42,47]. All methods were carried out in accordance with the European (2010/63/EU) and the ARRIVE guidelines. The experimental protocols, the animal handling, and the surgical and experimental procedures complied with the European guidelines (2010/63/EU) and Italian laws in force on the care and use of laboratory animals, and were approved by the Veterinarian Animal Care and Use Committee of the University of Parma and authorized by the Italian Ministry of Health.

#### Training and surgical procedures

The monkeys were first habituated to seat on a primate chair and to familiarize with the experimental setup. At the end of the habituation sessions, a head fixation system (Crist Instruments Co. Inc.) was implanted. Then, the monkeys were trained to perform the tasks described below. After completion of the training, a recording chamber (32×18 mm, Alpha Omega, Nazareth, Israel) was implanted on the ventrolateral prefrontal cortex (VLPF), based on MRI scan. All surgeries were carried out under general anesthesia (ketamine hydrocloride, 5 mg/kg, i.m. and medetomidine hydrocloride, 0.1 mg/kg, i.m.), followed by postsurgical pain medication.

### Experimental apparatus

During training and recording sessions, the monkeys seated on the monkey chair with the hand contralateral to the hemisphere to be recorded on a resting position, located 9 cm in front of the abdomen. A monitor was positioned in front of the monkey, to present the visual stimuli used in the passive tasks (Picture and Video tasks, see below). The monitor, with a resolution of 1680×1050 pixel, was positioned at 54 cm from monkey’s face, and its geometrical center was located at the height of monkey’s eyes. A laser spot could be projected on the center of the screen as fixation point. A phototransistor was placed on the monitor in order to provide the onset and offset of the visual stimuli.

During the Visuo-Motor task, a box containing three objects was positioned at 22 cm from the monkey’s chest. The opening of a small door (7×7 cm) in the frontal panel of the box at the height of monkey’s eyes allowed to present the three objects, one at the time. Two laser spots (instructing cues) of different colors (green and red) could be projected onto the box door or onto the object, signaling the task conditions and phases.

### Behavioral paradigms and stimuli

#### Picture task

The Picture task corresponds to the *Visual task* described in [42]. Briefly, to evaluate the response of VLPF neurons to the observation of static visual stimuli, 12 different images (6°x6°; see below) were presented, while the monkeys kept their gaze within a 6°x6° fixation window centered on the stimulus. Fig 1A shows the sequence of events occurring during each trial. The monkeys were required to keep their hand on the resting position; if this was accomplished, the trial started, and the fixation point (red laser spot) was turned on, and they had to fixate it for a randomized time interval (500-900 ms). If they kept fixation for this period of time, the fixation point turned off and one of the videos was presented for 600 ms, centered on the fixation point. The monkeys had to observe it (without breaking fixation) throughout the presentation period. Then, the image disappeared, the fixation point turned on again for a randomized period (500-900 ms) and the monkeys had to keep fixation on it.

The trials were accepted as correct, and the monkeys were rewarded, if they kept their eyes within the fixation window for the duration of each phase of the task and did not release the hand from the resting position. Discarded trials were repeated at the end of the sequence to collect at least 10 presentations for each stimulus. The order of stimuli presentation was randomized.

The 12 stimuli (Fig 1A) belong to 4 different semantic categories:

- Graspable solids (pictures of the objects employed in the motor task described in [38]: cube, cylinder, sphere);
- Fruits: apple, banana, peanut;
- Faces: human face, monkey face, sketchy drawing of a face;
- Laboratory furniture, geometric, but not graspable: shelf, monitor and clock.

The stimuli were homogeneous for luminance. Note that the fruits and solids could evoke similar affordances (apple and cube: power grip; banana and cylinder: finger prehension; sphere and peanut: precision grip).

#### Video Task

The Video corresponds to that described in [41]. In order to evaluate the response of VLPF neurons to observation of dynamic visual stimuli, we displayed videos (12°x12°) showing several biological stimuli and object motion (see below), while the monkey maintained fixation within a 6°x6° fixation window centered on the video. The sequence of events occurring during each trial is the same as in the Picture task (see Fig 1B), but the stimuli were presented for 1800 ms. The criteria employed for trial acceptance were the same as in the Picture task (see above).

Discarded trials were repeated at the end of the sequence in order to collect at least 10 presentations for each stimulus. The order of stimuli presentation was randomized.

The construction of the set of video stimuli was devised so to present to the monkey goal-related or non-goal-related actions, different agents and presence or absence of an object. Specifically, the 6 stimuli were the following (Fig 1B):

- Monkey grasping in first person perspective (*MGI):* a monkey right forelimb enters into the scene from the lower border of the video, reaches and grasps a piece of food located in the center of it, and lift it toward itself (only the initial phase of this latter movement is visible). The observed forelimb is presented as if the observing monkey was looking at its own forelimb during grasping.
- Monkey grasping in third person perspective (*MGIII):* a monkey, located in front of the observer, with its left forelimb reaches and grasps a piece of food located in the center of the video, and brings it toward itself.
- Human grasping (HG): a human actor, located on the right of the video reaches, grasps and lifts an object, located in the center of the video, with his right forelimb.
- Human mimicking (HM): a human actor, located on the right of the video, performs the pantomime of the same action shown in HG, without the object.
- Biological movement (BM): a human actor located on the right of the video extends his right forelimb, with the hand open, to reach the central part of the screen. No object is present.
- Object motion (OM): an object is presented in the center of the screen and moves along the same trajectory as in HG. This stimulus was obtained by removing the agent from HG, in order to have same stimulus kinematics as in HG.

In the videos the agents’ faces were not shown in order to avoid the possible influence of neural responses due to face presentation.

#### Visuo-Motor task

The Visuo-Motor task is the same described in [38,47]. Briefly, the task consisted of two basic conditions: Action and Inaction (Fig 1C). Each trial started with the monkeys’ hand on the starting position. Then, one of the two instructing cues (green=Action condition; red=Inaction condition) was turned on and projected onto a closed box door, placed in front of the monkeys. In both conditions, the monkeys had to maintain fixation within a 6°x6° fixation window centered on the instructing cue for a randomized time interval (500-1100 ms). Then, the box door opened allowing the monkeys to see one of three objects.

In the Action condition, during object presentation, the monkeys had to maintain fixation with the green cue still on, projected onto the object. After a randomized time (700 to 1100 ms), the green cue turned off (Go signal), instructing the monkeys to reach for, grasp the object and pull it.

In the Inaction condition, the trial unfolding and the events timing was the same as in the Action condition till the red cue turned off, after which the monkeys were required to keep fixation for a further 600 ms period, refraining from acting. The order of presentation of both, objects and conditions, was randomized.

If the monkeys correctly performed a trial, the reward was delivered at the end of it. A trial was discarded when one of the following types of error occurred: 1) releasing the hand from the resting position before reward delivery in the Inaction condition or before the Go signal in the Action condition; 2) breaking fixation before reward delivery in the Inaction condition or before the Go signal in the Action condition; 3) failing to reach and grasp the object; 4) grasping the object with an incorrect prehension. Discarded trials were repeated at the end of the sequence to collect at least 30 correct trials for condition (10 trials x 3 objects).

### Recording techniques, task events acquisition and microstimulation

Neuronal recordings were performed by means of a multi-electrode recording system (AlphaLab Pro, Alpha Omega Engineering, Nazareth, Israel) employing glass-coated microelectrodes (impedance, 0.5-1 MΩ) inserted through the intact dura. The microelectrodes were mounted on an electrode holder (MT, Microdriving Terminal, Alpha Omega) allowing electrodes displacement, controlled by a dedicated software (EPS; Alpha Omega). The MT holder was directly mounted on the recording chamber. Neuronal activity was filtered, amplified, and monitored with a multichannel processor and sorted using a multi-spike detector (MCP Plus 8 and ASD, Alpha Omega Engineering). Spike sorting was performed using the Off-line Sorter (Plexon, Inc, Dallas TX, USA). During each recording session, electrodes were inserted one after the other inside the dura until the first neuronal activity was detected for each of them. Each electrode was then deepened into the cortex independently one from the other, in steps of 500 µm (see [76] for a similar procedure). At each site, multiunit and single-unit activities were recorded for subsequent analyses. The experiment was controlled by a homemade Labview software. Digital output signals determined the onset and offset of laser spots, image/videoclip presentation, opening of the door and reward release. Contact-detecting electric circuits provided the digital signals related to monkey hand contact and release of the resting position and the beginning and end of object pulling.

In order to identify the sector where eye movements can be elicited by intracortical microstimulation, the recording microelectrodes were also used for delivering intracortical monophasic trains of cathodic square wave pulses, through a constant current isolator (World Precision Instruments, Stevenage, UK) with the following parameters: total train duration, 50 ms or, when no response was elicited, 100 ms; single pulse width, 0.2 ms; pulse frequency, 330 Hz. The stimulation started with a current intensity of 100 µA that was decreased until threshold definition and was controlled on an oscilloscope by measuring the voltage drop across a 10 KΩ resistor in series with the stimulating electrode.

Eye movements were recorded using an infrared pupil/corneal reflection tracking system (Iscan Inc., Cambridge, MA, USA) positioned above the box. Sampling rate was 120 Hz.

### Histology, reconstruction of the recorded area and identification of the regions of interest on the basis of architectural and connectional data

Before sacrificing the animals, electrolytic lesions (10 μA cathodic pulses per 10 s) were performed at known coordinates at the external borders of the recorded region. After one week, each animal was anaesthetized with ketamine cloride (15 mg⁄kg i.m.), followed by an i.v. lethal injection of pentobarbital sodium and perfused through the left cardiac ventricle with buffered saline, followed by fixative. The brain was then removed from the skull, photographed, frozen and cut coronally. Each second and fifth section (60 μm thick) of a series of five were stained using the Nissl method. The locations of penetrations were then reconstructed on the basis of electrolytic lesions, stereotaxic coordinates, depths of penetrations and functional properties. More specifically, penetrations deeper than 3000 μm located inside the Arcuate sulcus and Principal Sulcus were used in order to localize the posterior and medial border of VLPF, respectively.

In order to attribute the functional properties of each penetration to a specific VLPF area, we defined each anatomical area based on its cytoarchitectural and connectional properties. Specifically, the location of areas 46v, 12r, 45A, 45B, and 8 have been defined by analyzing the Nissl-stained sections of the recorded monkeys and by superimposing the average anatomical map of the VLPF described in [10]on their brain reconstructions. In addition, since in the previous studies of our lab [11,29,31], areas 12r and 46v have been further subdivided in three parts, based on their connectional properties, an additional procedure has been performed. In particular, to identify these parts, a non-linear transformation procedure has been used (for details see [46] to superimpose the location of the injection sites identified in our previous works on the histological reconstruction of the two recorded brains, based on specific anatomical anchor points: anterior and posterior tip of the principal sulcus, spur of the arcuate sulcus, tip of the superior and inferior arcuate sulcus, tips of the superior and inferior prefrontal dimples, orbital reflection at the lateral orbital sulcus level. This approach allowed us to further subdivide areas 46v and 12r, and to confirm the architectural data used for identifying areas 45A, 45B, and 8. In this work, for sake of simplicity, we refer to anatomical areas caudal 46vc, rostral 46vc, intermediate 12r and 46vr as caudal 46v, middle 46v, middle 12r and rostral 46v, respectively.

In addition, we defined an oculomotor prearcuate sector by using microstimulation and recording of saccade-related neuronal activity. More in details, concerning microstimulation, oculomotor sites were those in which saccadic eye movements could be elicited with currents lower of or equal to 60 µA, with a train duration of 50 or 100 ms; concerning single neuron recordings, we described saccade-related activity by aligning the neuronal discharge with the moment in which the eyes reached a fixation target. This region, overlapping the location of areas 8 and 45B, has been excluded from further analyses.

The location of our recording chamber allowed us to record only from a small number of sites falling in areas rostral 46v and rostral 12r, especially in M2, thus, we pulled together the data obtained from these areas and named this region *rostral sector*.

### Analysis of single neurons responses

The digital signals representing the different task events, described above, were employed to align neuronal activity and to create the response histograms and data files used for the statistical analyses described in the subsequent paragraphs.

#### Picture task

We recorded neuronal activity for at least 120 successful trials, 10 for each stimulus. For the statistical analysis, two epochs were defined (see 42): 1) *Baseline*: 500 ms preceding stimulus presentation, during which the monkey was looking at the fixation point; *2*) *Presentation*: the first 500 ms of image presentation.

Single-neuron responses were statistically evaluated by means of a 2X12 ANOVA (Factors: Epoch, Stimulus, p<0.01) followed by Newman-Keuls post hoc tests.

A neuron was considered as task-related when the 2X12 ANOVA revealed: 1) a significant main effect of the Epoch factor (p<0.01) and/or 2) a significant interaction effect (Epoch x Stimulus, p<0.01), with the post-hoc tests showing a significant difference between at least one *Presentation* epoch of one image and its corresponding *Baseline epoch*. Task-related neurons were classified as *selective* when the 2X12 ANOVA revealed a significant Interaction effect and the post-hoc test showed a significant difference among the activity recorded in the *Presentation epoch* of one image and that of its corresponding *Baseline epoch* as well as a significant difference between the activity recorded in the *Presentation epoch* of that image and the *Presentation epoch* of at least another image. Neurons were classified as *unselective* when the statistical test revealed a significant Main effect of the Epoch factor and/or a significant Interaction effect, and the post-hoc test did not show any difference among the activities recorded in the *Presentation* epochs of the 12 images.

#### Video task

We recorded neuronal activity for at least 60 successful trials. For the statistical analysis, three epochs were defined (see 41): 1) *Baseline*: 500 ms before the beginning of the videos, during which the monkey was looking at the fixation point; 2) *Video Epoch 1*: the first 700 ms of the videos (except for MGI, where, because of the fastest arm movement, the epoch lasted 500 ms); 3) *Video Epoch 2*: the subsequent 700 ms of the videos. Note that Epoch 1 includes the context of the scene and the beginning of the forelimb movement, while Epoch 2 includes the hand-object contact in the case of action and the end of movement/object motion in all other videos.

Single-neuron responses were statistically evaluated by means of a 3X6 ANOVA (Factors: Epoch, Stimulus, p<0.01) followed by Newman–Keuls post hoc tests.

A neuron was considered as task-related when the 3X6 ANOVA revealed: 1) a significant main effect of the Epoch factor (p<0.01), with post-hoc tests indicating a significant difference between at least one of the two *video epochs* and the baseline and/or 2) a significant interaction effect (Epoch x Stimulus, p<0.01), with post-hoc tests showing a significant difference between at least one of the *video epochs* of one video and the corresponding *baseline* epoch.

Task-related neurons were classified as *selective* when the 3X6 ANOVA revealed a significant Interaction effect and the post-hoc test showed a significant difference among the activity recorded in one of the two *Video epoc*hs of one video and that of the corresponding *Baseline epoch* as well as a significant difference between the activity recorded in one of the two *Video epochs* and that recorded in the same epochs of at least another video. Neurons were classified as *unselective* when the statistical test revealed a significant Main effect of the Epoch factor and/or a significant Interaction effect, and the post-hoc test did not show any difference among the activities recorded in the *Video epochs* of the 6 videos.

#### Visuo-Motor task

We recorded neuronal activity for at least 60 successful trials (thirty per condition, 10 for each object). For statistical analysis of the neural activity, nine epochs have been defined (see *38, 42*), based on the digital signals: 1) Baseline: from 750 ms to 250 ms before the onset of the instructing cue; 2) Pre-cue: 250 ms preceding the onset of the instructing cue; 3) Cue: 250 ms following the onset of the instructing cue; 4) Pre-presentation: 500 ms preceding the opening of the box door; 5) Presentation: 500 ms following door opening (object presentation); 6) Set: 250 ms before the offset of the instructing cue; 7) Go/NoGo, from the offset of the instructing cue to the release of the hand starting position (Action condition) or 250 ms following the offset of the instructing cue (Inaction condition); 8) Grasping/Fixation: from 250 ms before to 250 ms after the Pulling onset (Action condition) or a time period ranging from 250 ms to 500 ms after the offset of the instructing cue (Inaction condition); 9) Reward: 500 ms following reward delivery.

Single-neuron responses were statistically evaluated by means of a 9X2 ANOVA (Factors: Epoch, Condition, p<0.01) followed by Newman-Keuls post hoc tests. Since trials were randomized, changes of the baseline activity across trials were not expected, and the neurons showing a significant difference between baselines were discarded. Neurons were included in our dataset and were defined as *task related* when the 9×2 ANOVA revealed at least one of the two following effects: 1) a significant main effect of the Epoch factor (p<0.01), with the relative post-hoc tests showing a significant difference between the activity recorded in the Baseline epoch and in at least one of the other epochs (Condition-independent neurons); 2) a significant Interaction effect (Condition x Epoch, p<0.01), with the subsequent post-hoc tests showing a significant difference between at least one epoch of one condition and both its baseline and the corresponding epoch of the other condition (Condition-dependent neurons). Considering that the epochs of Pre-cue and Reward fall in the inter-trial period, when eye movements are not controlled, we decided to consider, for our analysis, the remaining six epochs plus the Baseline.

### Population analyses

Since in all the tasks considered in this work most time intervals were randomized, similarly to single neuron analysis, for each task, we considered a time period of 6 s centered on each task event and joined them in a matrix of the same time length for each trial.

In the subsequent analyses (time course of mean population activity, demixed principal component and decoding analyses) we employed different time periods extracted from this matrix.

To characterize the time course and the discharge rate of different neuronal populations with respect to the main tasks phases, the neuronal activity of each population was aligned with the main behavioral events. The population activity was computed as follows. The mean single neuron activity over trials, in terms of firing rate, was calculated for each 20 ms bin in the two conditions. The average baseline activity was then subtracted from the mean single neuron activity over trials for each bin. Thus, in this analysis, 0 represents baseline activity. The net average discharge frequency of each neuron was used for subsequent statistical analysis. Each neuron contributed one entry to each data set.

#### Demixed principal component analysis

In order to evaluate how different populations of neurons encode specific factors during the unfolding of the Visuo-Motor task, we adopted a data-simplification method: the demixed principal component analysis (dPCA), using freely available code provided by Kobak and coworkers ([77],see also [47]).

dPCA is a supervised dimensionality reduction method which allows to account for information related to the specific task factors. In this work, we considered the Type of Stimulus factor for the Picture and Video tasks (including the 12 stimuli and 6 videos, respectively), and the Condition (Action and Inaction) and Object (Cube, Cylinder and Sphere) factors for the Visuo-Motor task. This approach allowed to extract the demixed principal components explaining variance which is mainly related to the considered factors as well as variance unrelated to the chosen factors. These components were thus used to obtain an isolated description of the respective neural modulations.

In addition, the toolbox allows to obtain an estimate of the signal variance, computed after removing the estimated noise-related variance, which is later partitioned into signal variance that is related or unrelated to the chosen factors. To obtain a more reliable representation of each of these partitions of the signal variance, we computed a weighted value by multiplying them to the total signal variance observed for each area and period analyzed, and then dividing the results by 100. This weighted value is the one reported in the results section.

For the Picture and Video tasks, we considered a Presentation period corresponding to 500 and 1400 ms following stimulus appearance, respectively.

For the Visuo-Motor task, we used the following time periods: Cue period: 500 ms following cue onset; Presentation period: 500 ms following object appearance; Go/NoGo-Behavioral response period, formed, in the Action condition, by merging the 200 ms following the Go signal and the subsequent 400 ms, starting 300 ms before and ending 100 ms after the beginning of object holding; in the Inaction condition by the 600 ms following the NoGO signal. Then, we plotted the time course of the first principal component related to each of the considered factors. The toolbox uses a linear classifier (stratified Monte Carlo leave-group-out cross validation) to evaluate at which time points the given task elements belonging to a factor (i.e. Type of Stimulus in the Picture and Video tasks; Condition and Object in the Visuo-Motor task) are significantly different from each other. As criterion for significant statistical separation of the curves, the actual classification accuracy had to be at least 3 standard deviations above the mean chance accuracy, obtained after 100 shuffled repetitions of the analysis, considering a time period of at least 5 consecutive 20 ms bins.

#### Decoding analysis

In order to estimate which type of information is coded by the different neuronal populations recorded in the Visuo-Motor task and how this information is encoded in static or dynamic patterns of activity, we adopted a population decoding approach according to the methodology described by Meyers and coworkers [78,79].

Based on the above defined population matrix, we identified the following time periods: a 500 ms Baseline period, starting 750 ms before cue onset; a Cue period of 500 ms following cue onset; a Presentation period of 500 ms following object presentation; a Decision period of 400 ms starting 200 ms before the Go/NoGo signal; a Behavioral response period of 400 ms, starting, in the Action condition, 300 ms before object holding, and in the Inaction condition, 200 ms after the NoGo signal.

Decoding analysis was performed as described in [47]. In order to evaluate the temporal evolution of information coding, we applied a cross-temporal decoding analysis, which consists in training the classifier at time t and testing it at the same time t (on-diagonal bins) and at all the other times (off-diagonal bins). With respect to the results of cross-temporal decoding, our definition of *static patterns of coding* refers to the situations in which off-diagonal bins are associated to high accuracy values [79].

The data alignment on task events and the binning procedure described above led to merge in the same bin the activity at the border between two subsequent periods of the task (bins of 60 ms, sampled at 20 ms intervals). Accordingly, in our analysis, we removed the last two bins of each task period considered in the analysis, obtaining a total number of 105 considered time bins.

#### Classification of static patterns

To investigate whether the observed *static patterns of coding* are statistically significant, we used a method employed in previous studies [79,80]. First, for each off-diagonal bin, we calculated the differences between its accuracy value and that of the two on-diagonal bins used for training and testing. The same procedure was repeated 1000 times using accuracies obtained after shuffling the labels, allowing us to obtain a null distribution of the differences in accuracy values between the off- and on-diagonal bins. For statistical analysis we selected those time bins in which both differences were lower than 99.9% of the differences estimated at the corresponding time points in the null distribution. Then, we identified clusters of consecutive off-diagonal time bins (n. bins > 0) associated to differences exceeding the 99.9% critical value, and summed the differences observed in each point of the cluster. The same procedure was applied using the accuracy value matrices resulting from the 1000 shuffled decodings, each time extracting the maximum summed cluster values.

Finally, we performed a cluster-based permutation test [81], by comparing to the summed difference values of the observed-data clusters with the maximum summed cluster values of the null distribution. The proportion (over the 1000 iterations) of times in which the maximum summed cluster values of the null distribution were lower than the values obtained in each observed-data cluster determined the p-value of the test. The bins for which this p value was below 0.001 were considered significantly static off-diagonal bins. Similar to [79] as a further restrictive criterion to classify the off-diagonal bins as significantly static, the two corresponding on-diagonal bins used for training and testing the classifier were required to be both significantly above chance level. In this case, we performed a permutation test consisting in evaluating the proportion of times in which the accuracy value observed in each on-diagonal time bin was higher than that observed in the null distribution, which determined the p-value of the test. The accuracy in an on-diagonal time bin was then considered significantly above chance if the obtained p-value was below < 0.000009, corresponding to a p-value of 0.001 Bonferroni corrected for the 105 considered bins.

## Acknowledgments

We thank M. Bimbi for his technical help in setting up the experiment.

## Supporting information

**S1 Fig.**
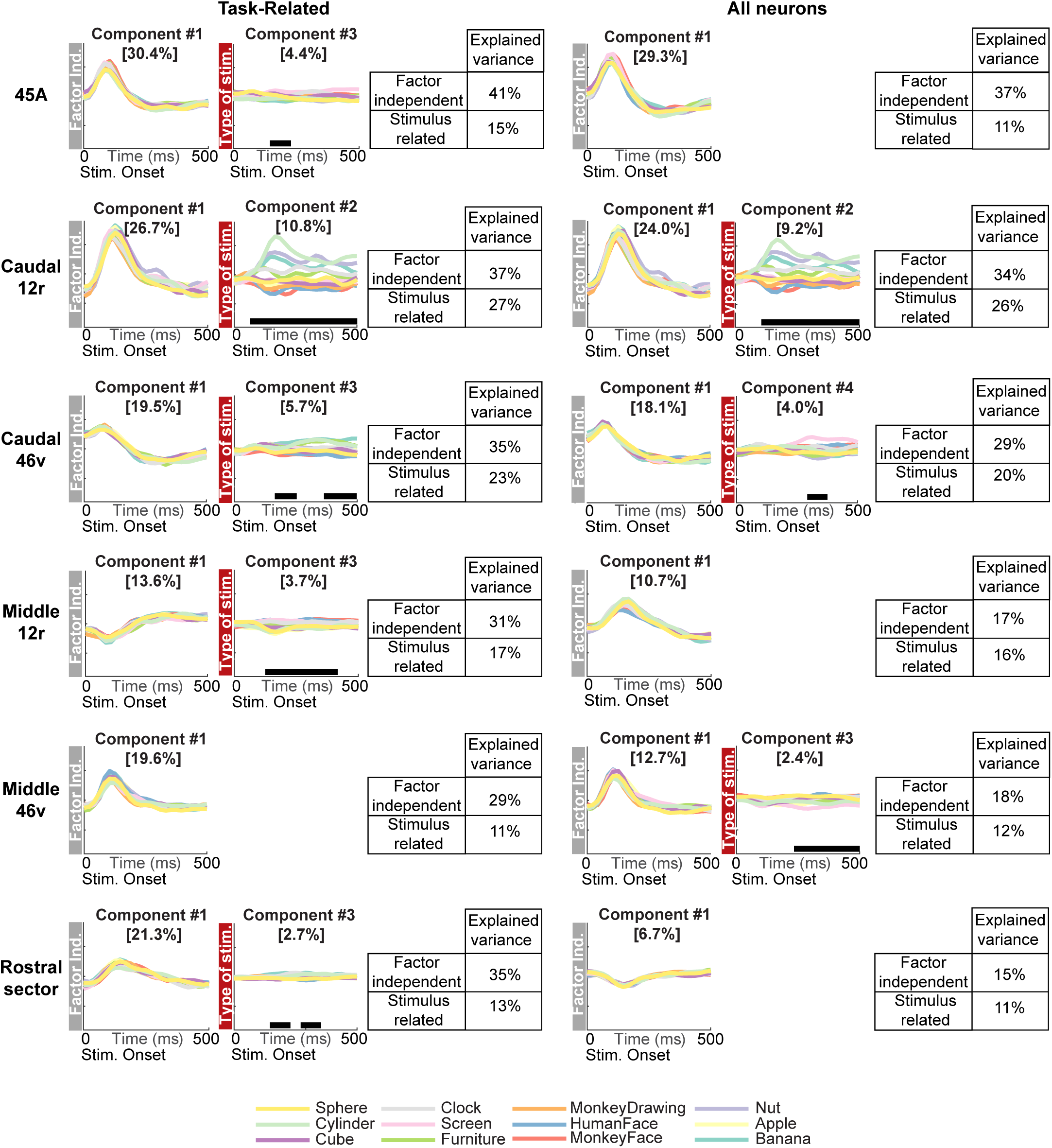
dPCA of the population activity of neurons recorded in the Picture task. Results of dPCA performed on the Presentation period of the Picture task (see Methods), considering the task-related neurons (left column) and the whole population of recorded neurons (right column). Each panel depicts the time course and the relative percent of explained variance of the first Factor-Independent principal component and of the first significant Type of Stimulus-related principal component. Each colored line corresponds to one of the twelve stimuli presented in the task. Horizontal thick lines indicate the time intervals where the task factor is significantly decoded (see Methods). For each area, the table on the right shows the Factor-independent and Type of Stimulus percentage of total variance.

**S2 Fig.**
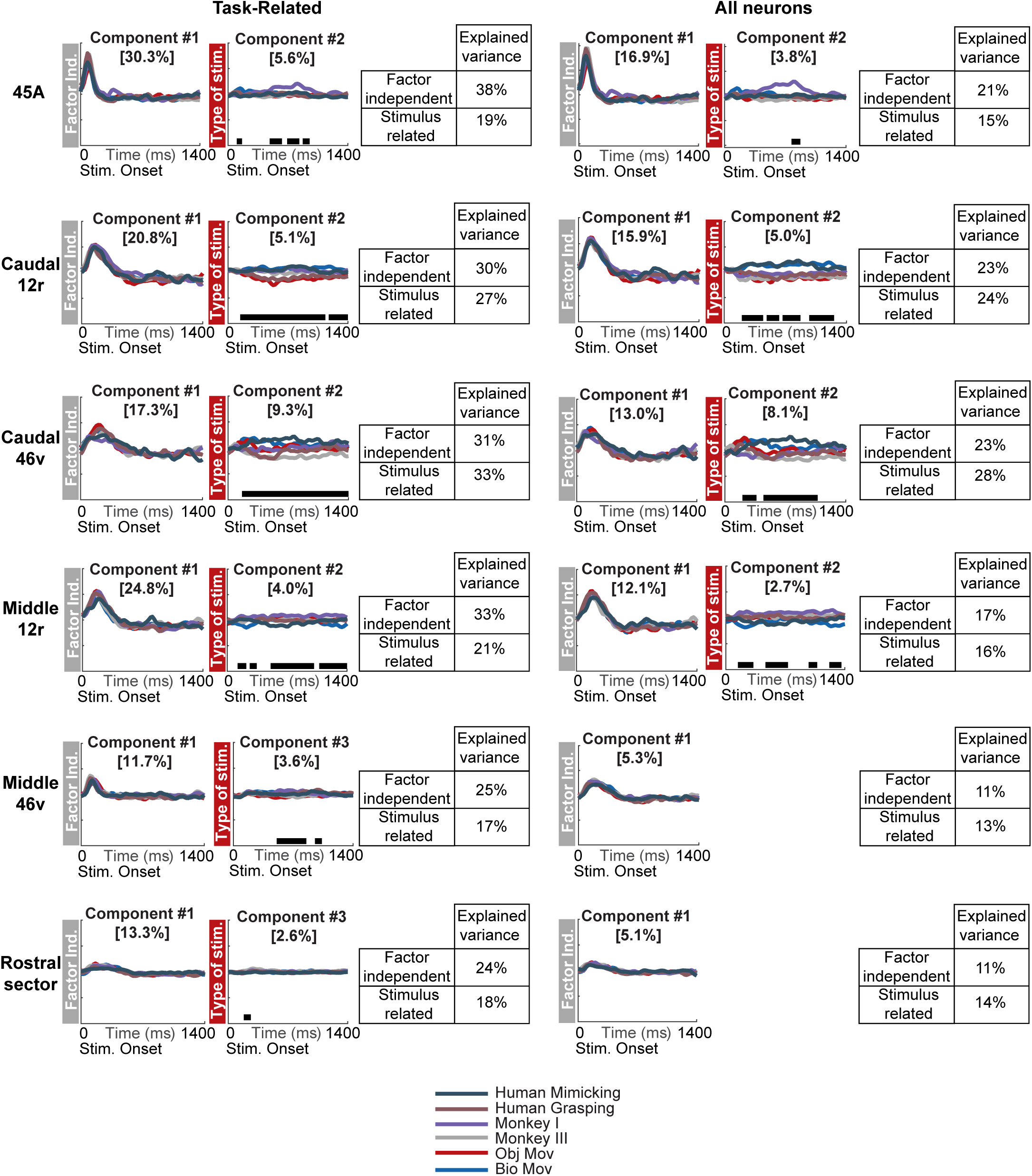
dPCA of the population activity of neurons recorded in the Video task. Results of dPCA performed on the Presentation period of the Video task (see Methods), considering task-related neurons (left column) and whole population of recorded neurons (right column). Each panel depicts the time course and the relative percent of explained variance of the first Factor-Independent principal component and of the first significant Type of Stimulus-related principal component. Each colored line corresponds to one of the six stimuli presented in the task. Horizontal thick lines indicate the time intervals where the task factor is significantly decoded (see Methods). For each area, the table on the right shows the Factor-independent and Type of Stimulus percentage of total variance.

**S3 Fig.**
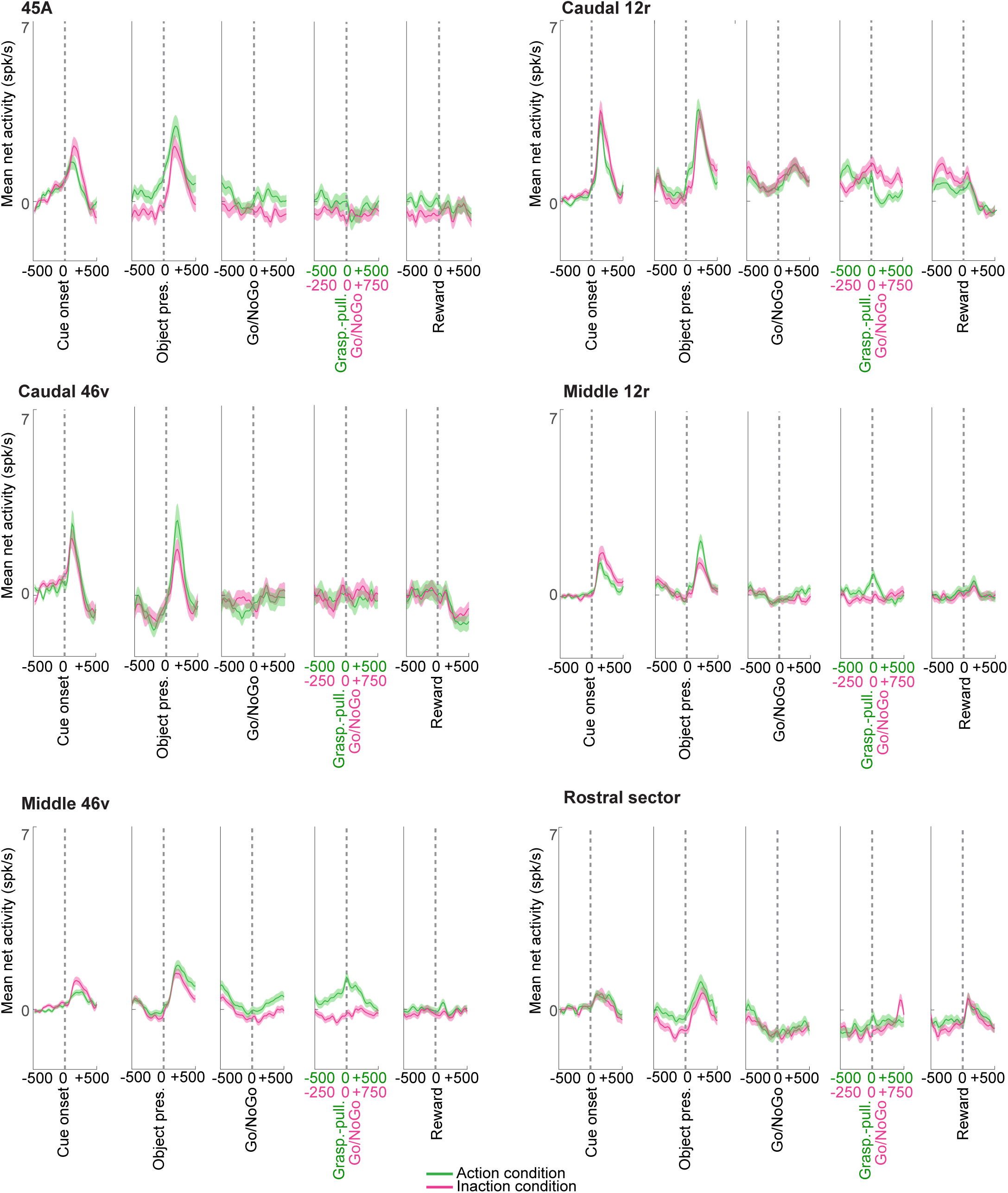
Mean activity of the whole population of neurons recorded in each area during the Visuo-Motor task. Temporal profile of the mean net activity of the whole population of recorded neurons (including both task-related and non-task-related neurons). Magenta and green curves indicate the population mean net activity in the Inaction and Action condition, respectively. The shaded area around each curve represents standard error. The neuronal activity is aligned on the main task events indicated below each panel.

**S4 Fig.**
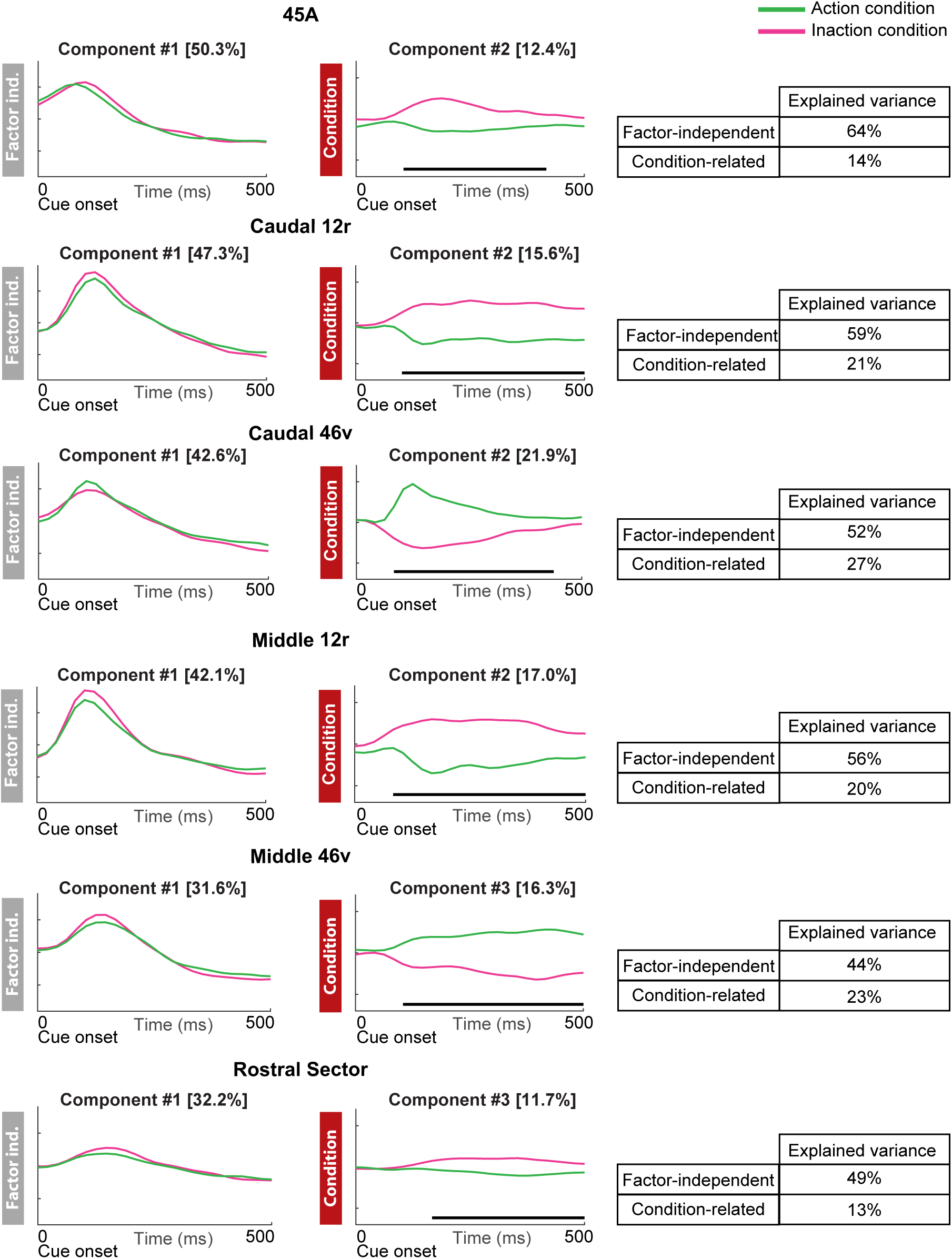
dPCA of the population activity of task-related neurons recorded during the Cue period of the Visuo-Motor task. Each panel depicts the time course and the percent of explained variance of the first Factor-Independent principal component and of the first significant Condition-related principal component. Horizontal thick lines indicate the time intervals where the task factor is significantly decoded (see Methods). For each area, the table on the right shows the Factor-independent and Condition-related percentage of the total variance.

**S5 Fig.**
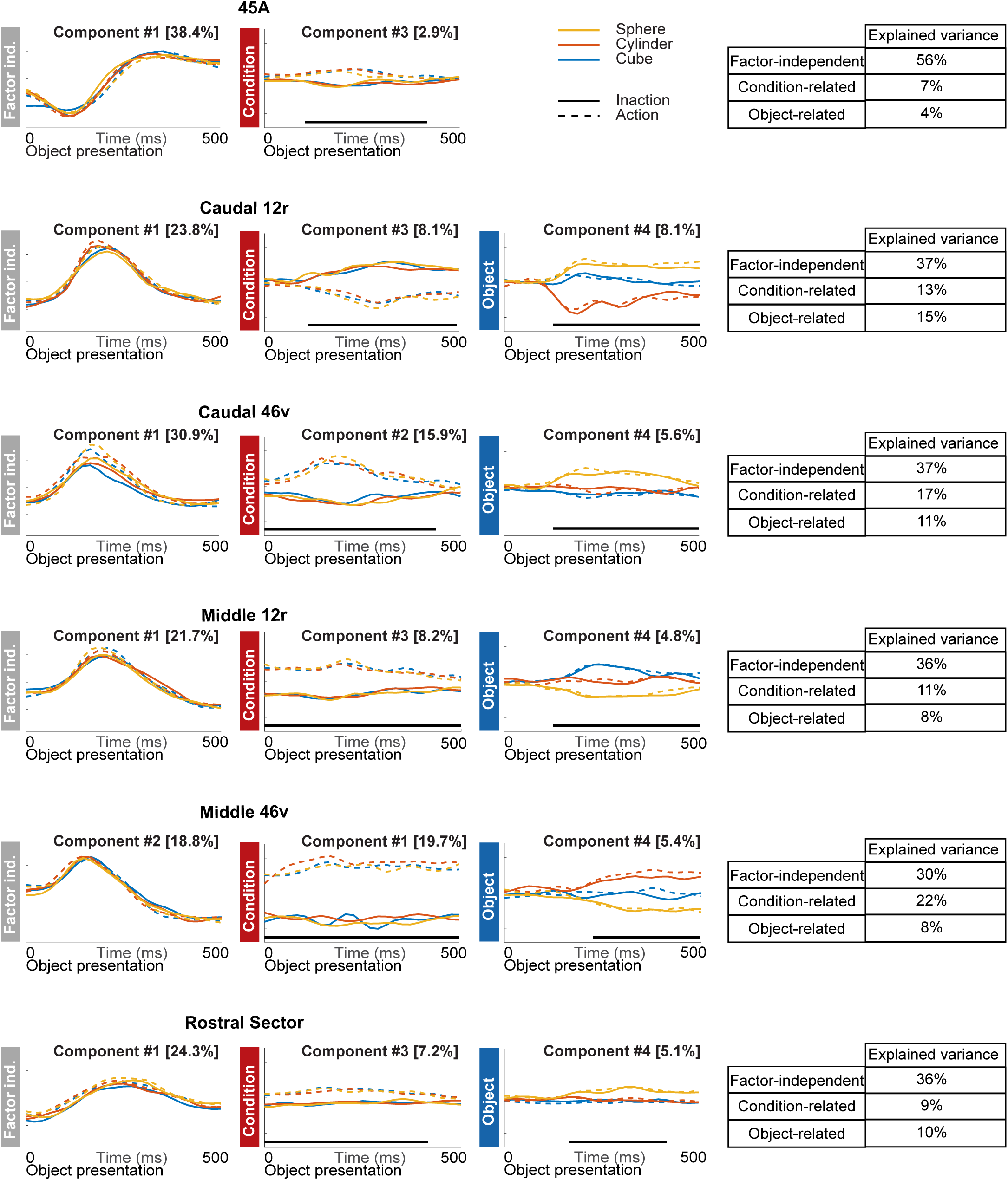
dPCA of the population activity of task-related neurons recorded during the Presentation period of the Visuo-Motor task. Each panel depicts the time course and the percent of explained variance of the first Factor-Independent principal component and of the first significant Condition/object-related principal components. Horizontal thick lines indicate the time intervals where the task factor is significantly decoded (see Methods). For each area, the table on the right shows the Factor-independent, Condition-related and Object-related percentage of the total variance.

**S6 Fig.**
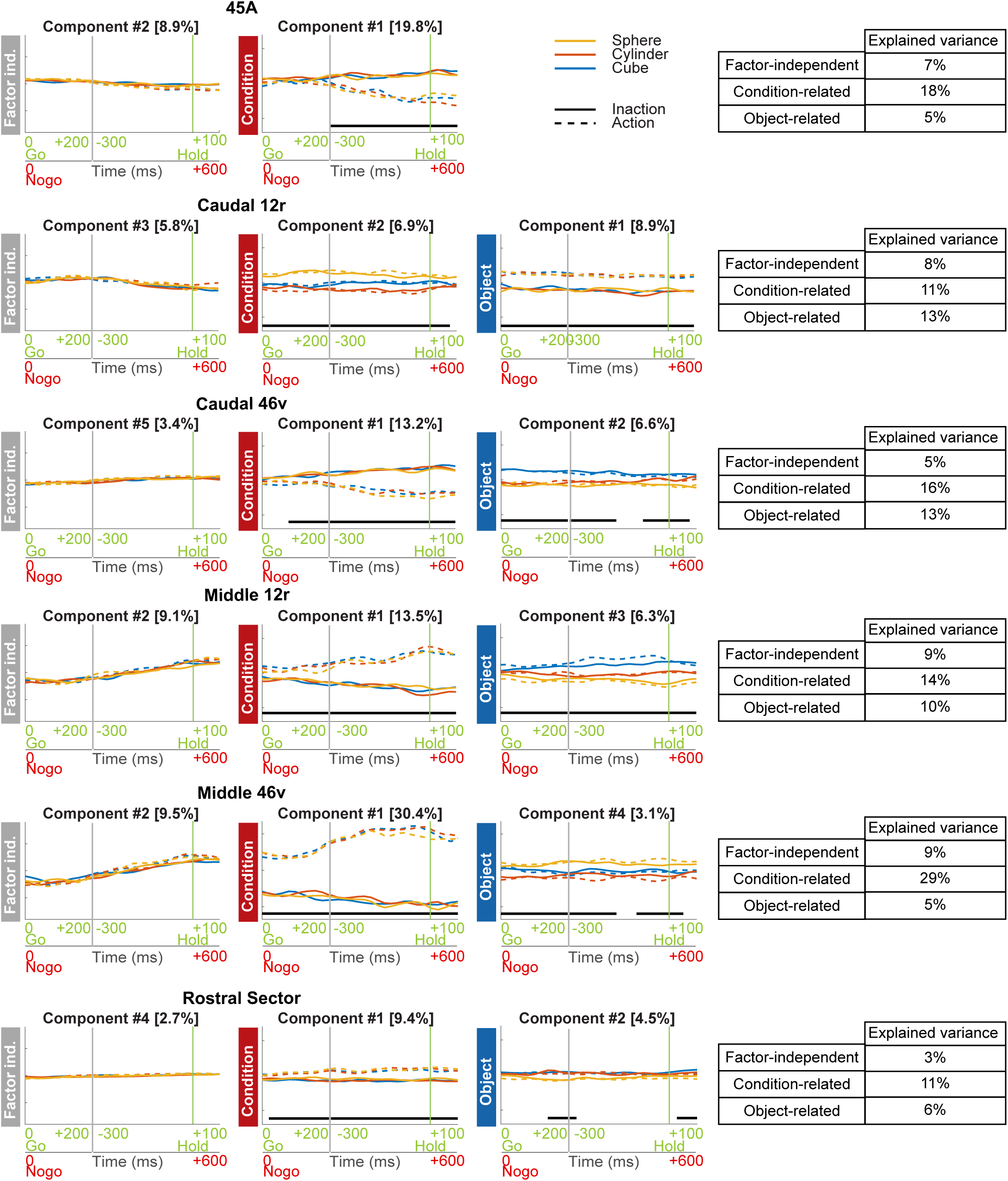
dPCA of the population activity of task-related neurons recorded during the Go/NoGo-Behavioral response period of the Visuo-Motor task. Each panel depicts the time course and the percent of explained variance of the first Factor-Independent principal component and of the first significant Condition/object-related principal components. Horizontal thick lines indicate the time intervals where the task factor is significantly decoded (see Methods). The vertical grey line separates Go/NoGo and Behavioral response periods (see Methods). The vertical green line indicates the beginning of holding period. For each area, the table on the right shows the Factor-independent, Condition-related and Object-related percentage of the total variance.

**S7 Fig.**
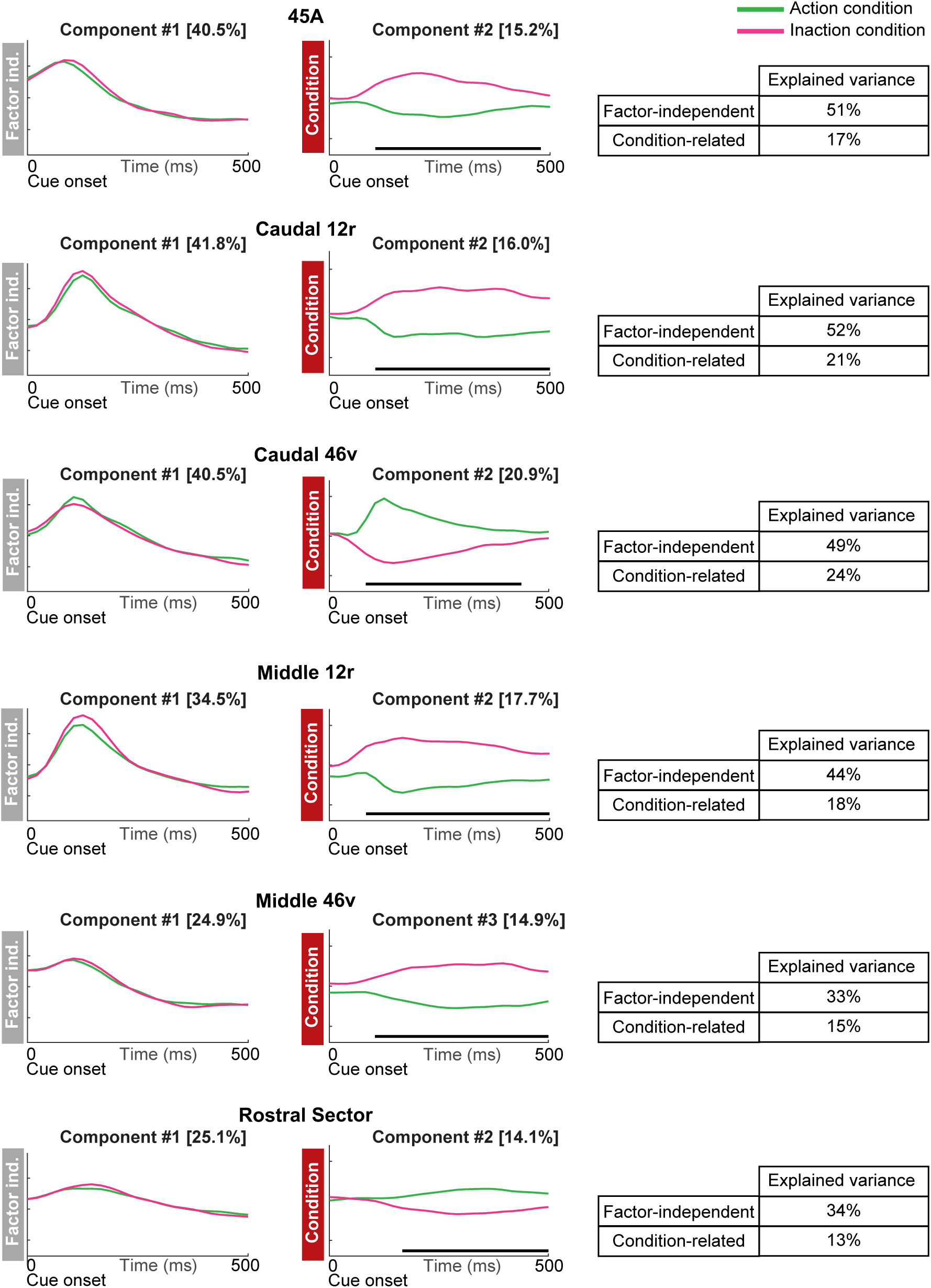
dPCA of the activity of the whole population of neurons recorded during the Cue Period of the Visuo-Motor task. Each panel depicts the time course and the percent of explained variance of the first Factor-Independent principal component and of the first significant Condition-related principal component. Horizontal thick lines indicate the time intervals where the task factor is significantly decoded (see Methods). For each area, the table on the right shows the Factor-independent and Condition-related percentage of the total variance.

**S8 Fig.**
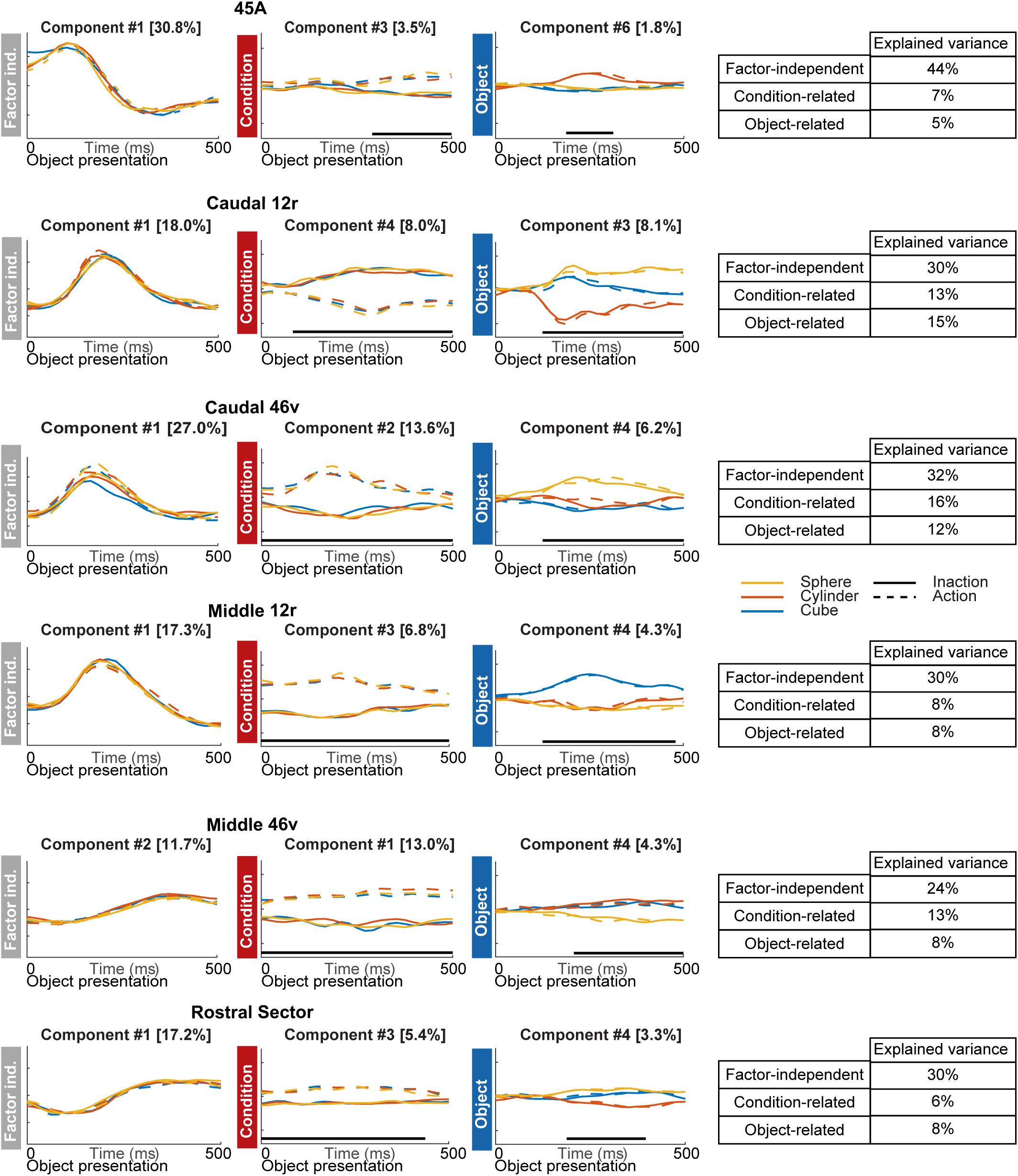
dPCA of the activity of the whole population of neurons recorded during the Presentation Period of the Visuo-Motor task. Each panel depicts the time course and the percent of explained variance of the first Factor-Independent principal component and of the first significant Condition/object-related principal components. Horizontal thick lines indicate the time intervals where the task factor is significantly decoded (see Methods). For each area, the table on the right shows the Factor-independent, Condition-related and Object-related percentage of the total variance.

**S9 Fig.**
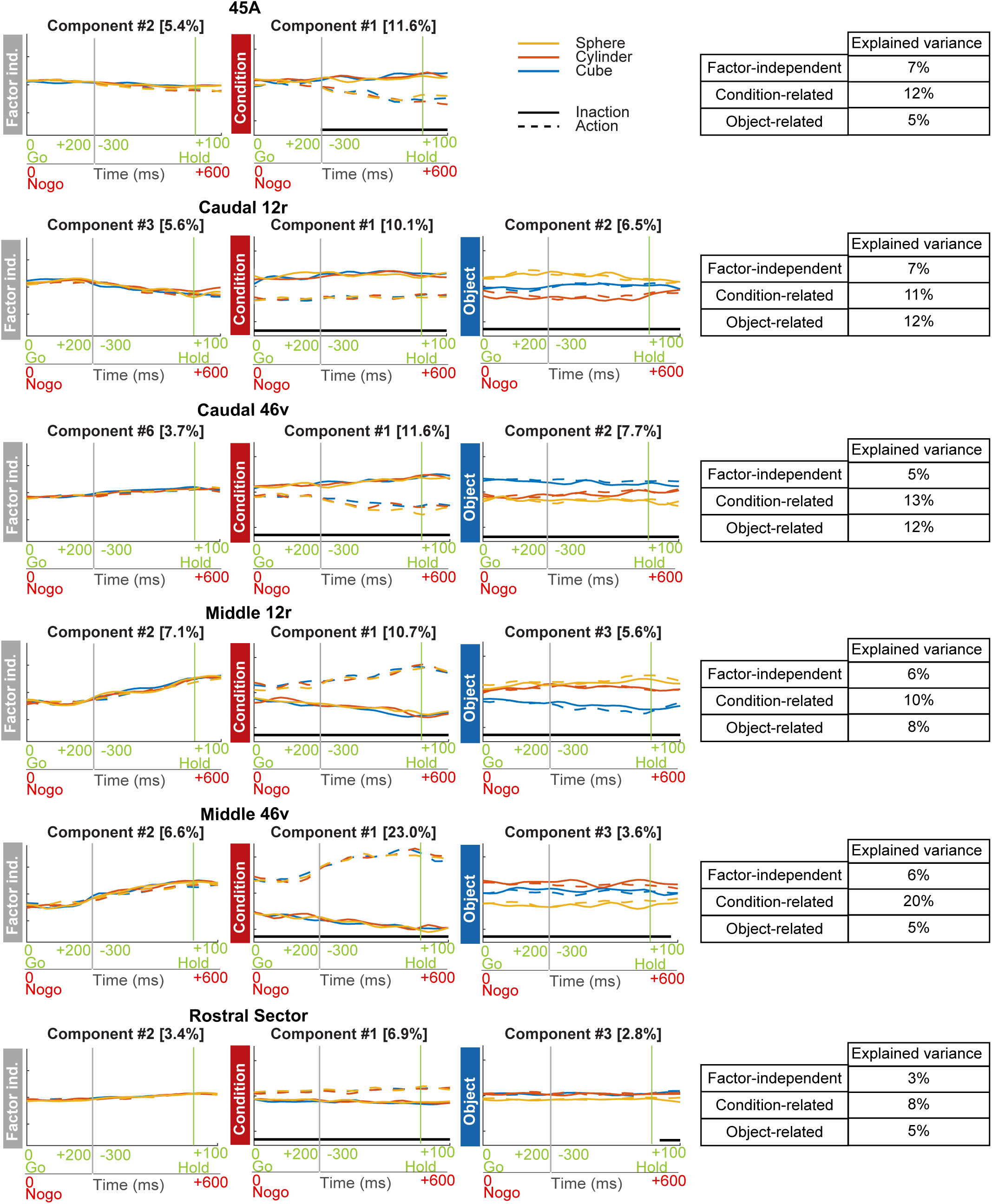
dPCA of the activity of the whole population of neurons recorded during the Go/NoGo-Behavioral response Period of the Visuo-Motor task. Each panel depicts the time course and the percent of explained variance of the first Factor-Independent principal component and of the first significant Condition/object-related principal components. Horizontal thick lines indicate the time intervals where the task factor is significantly decoded (see Methods). The vertical grey line separates Go/NoGo and Behavioral response periods (see Methods). The vertical green line indicates the beginning of holding period. For each area, the table on the right shows the Factor-independent, Condition-related and Object-related percentage of the total variance.

**S10 Fig.**
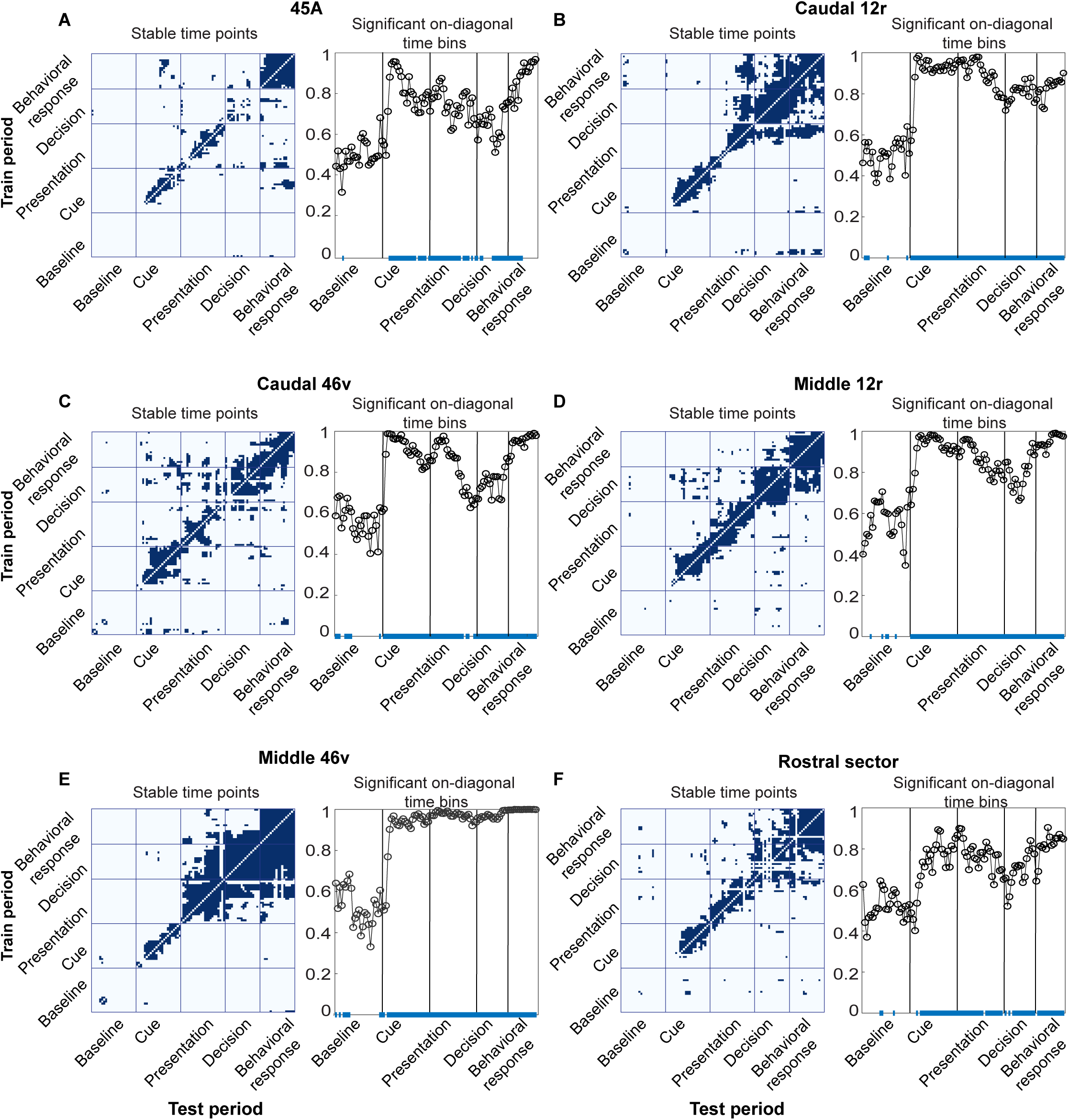
Significance of Condition decoding time points. Results are related to the decoding analysis presented in the main text and performed on the task related neurons recorded in each area. For each VLPF area, the left panel depicts the time points that have been classified as stable after cross-temporal Condition decoding (see Methods); the right panel depicts the accuracy levels of the Condition decoding observed along the diagonal during task unfolding (see Methods). The blue line on the bottom of the accuracy plots represents on-diagonal time points in which the accuracy level is significantly above chance (see Methods).

**S11 Fig.**
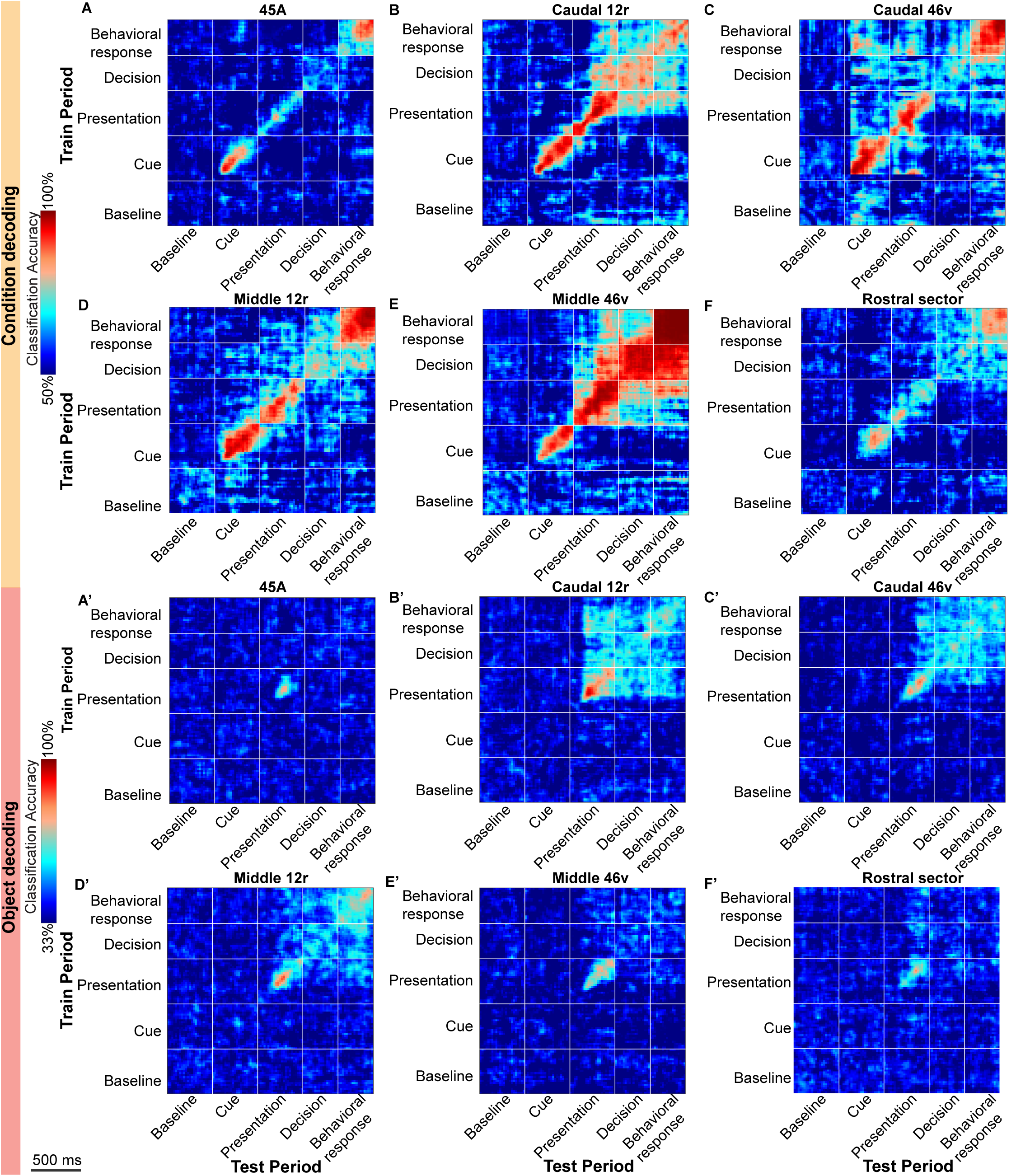
Significance of Object decoding time points. Results are related to the decoding analysis presented in the main text and performed on the task related neurons recorded in each area. For each VLPF area, the left panel depicts the time points that have been classified as stable after cross-temporal Object decoding (see Methods); the right panel depicts the accuracy levels of the Object decoding observed along the diagonal during task unfolding (see Methods). The blue line on the bottom of the accuracy plots represents on-diagonal time points in which the accuracy level is significantly above chance (see Methods).

**S12 Fig.**
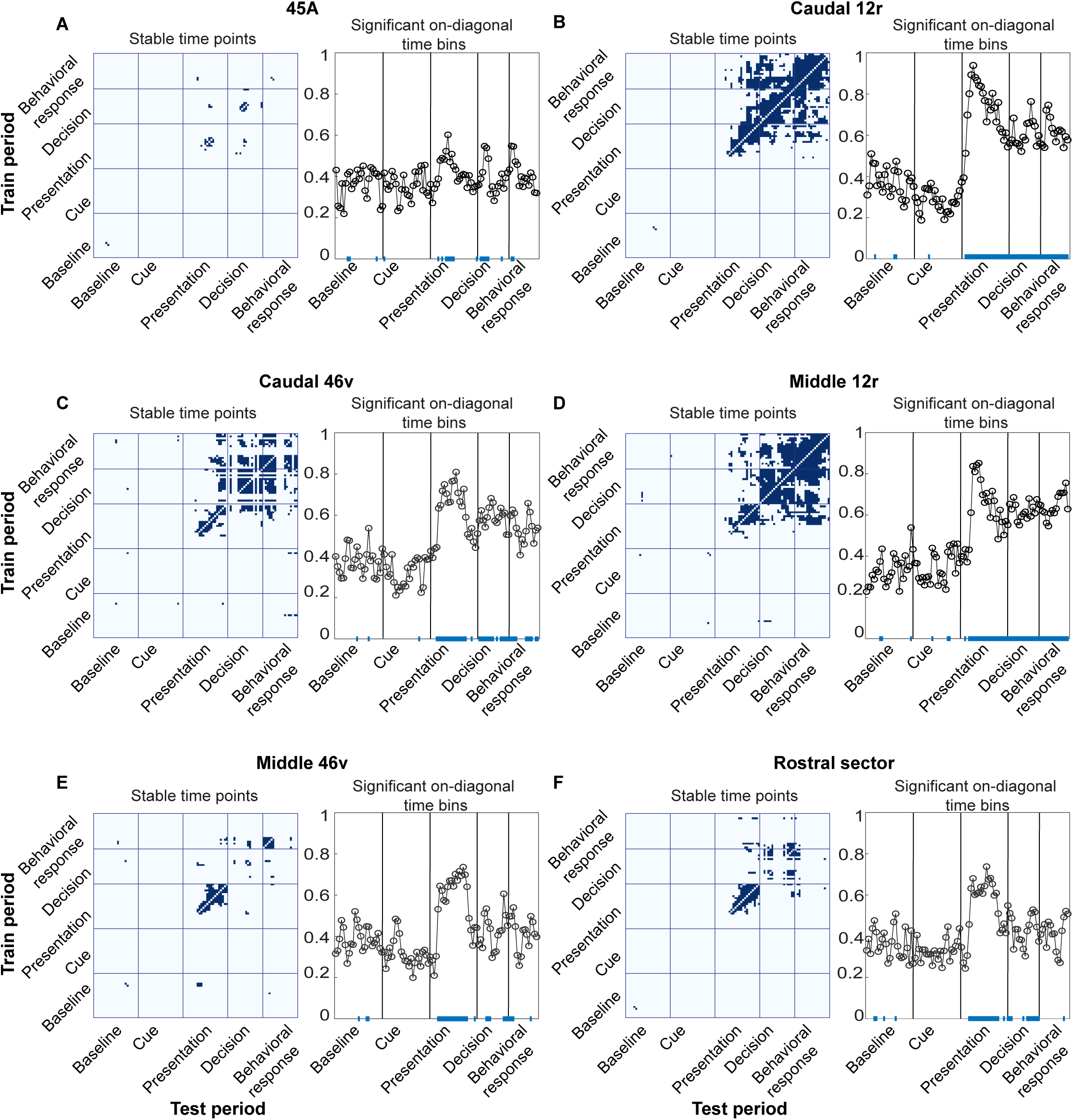
Cross-temporal decoding of the Condition (A-F) and Object (A’-F’) factors of the Visuo-Motor task in the whole population of recorded neurons. For each analysis, the decoding accuracy is computed in bins of 60 ms, sampled at 20 ms intervals. For each plot, the vertical and horizontal lines delimit the considered time periods (see Methods). Decoding periods of testing and training are indicated on the X and Y axes, respectively.

**S13 Fig.**
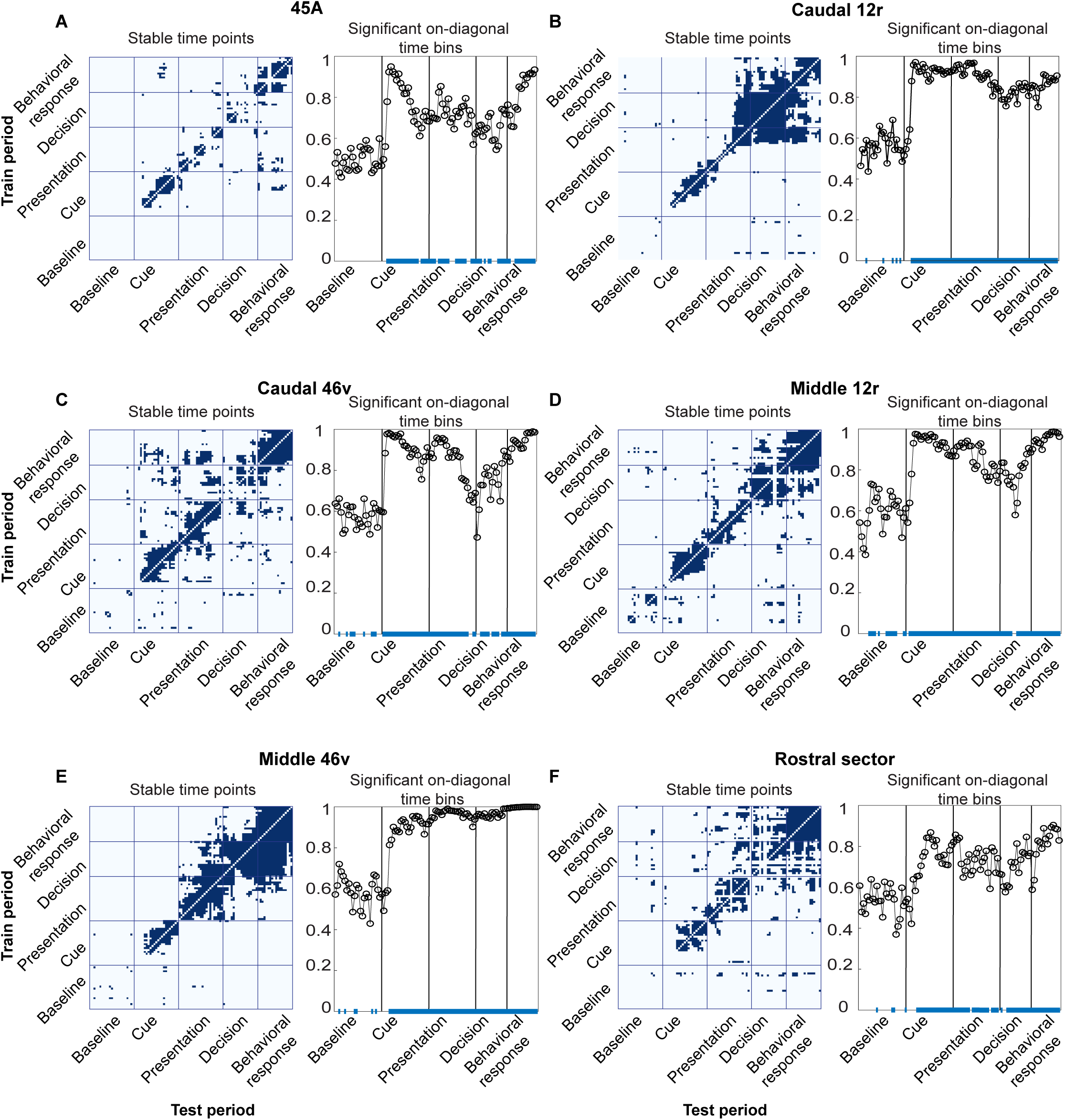
Significance of Condition decoding time points. Results are related to the decoding analysis performed on the whole population of neurons recorded in each area. For each VLPF area the left panel depicts the time points that have been classified as stable after cross-temporal Condition decoding (see Methods); the right panel depicts the accuracy levels of the Condition decoding observed along the diagonal during task unfolding (see Methods). The blue line on the bottom of the accuracy plots represents on-diagonal time points in which the accuracy level is significantly above chance (see Methods).

**S14 Fig.**
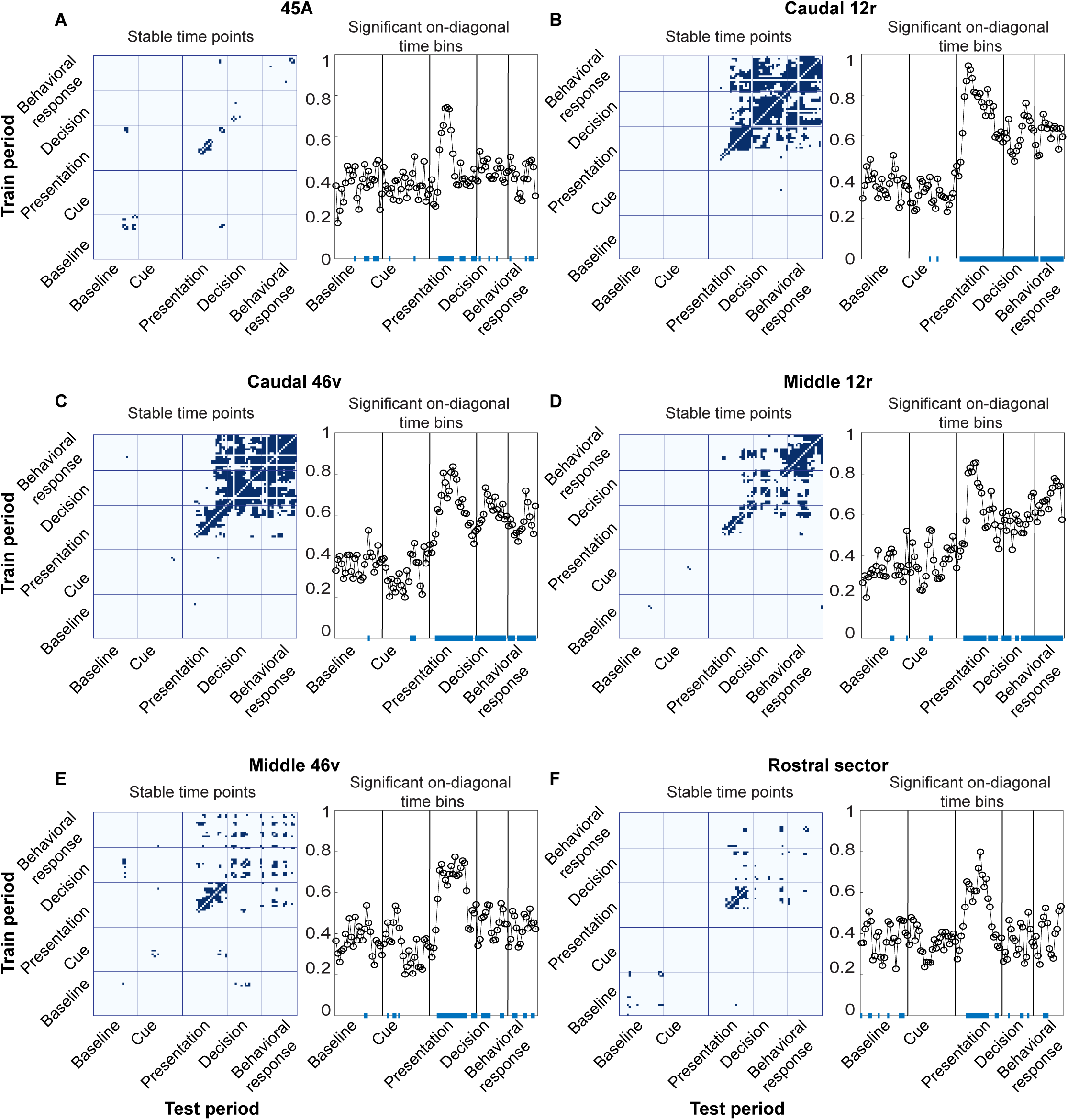
Significance of Object decoding time points. Results are related to the decoding analysis performed on the whole population of neurons recorded in each area. For each VLPF area the left panel depicts the time points that have been classified as stable after cross-temporal Object decoding (see Methods); the right panel depicts the accuracy levels of the Object decoding observed along the diagonal during task unfolding (see Methods). The blue line on the bottom of the accuracy plots represents on-diagonal time points in which the accuracy level is significantly above chance (see Methods).

## References

1. Levy R. The prefrontal cortex: from monkey to man. Brain. 2024;147: 794–815. doi:10.1093/brain/awad389

2. Miller EK, Cohen JD. An Integrative Theory of Prefrontal Cortex Function. Annu Rev Neurosci. 2001;24: 167–202. doi:10.1146/annurev.neuro.24.1.167

3. Fuster JM. Neurophysiology. The Prefrontal Cortex. Elsevier; 2015. pp. 237–308. doi:10.1016/B978-0-12-407815-4.00006-4

4. Asaad WF, Rainer G, Miller EK. Neural activity in the primate prefrontal cortex during associative learning. Neuron. 1998;21: 1399–407. doi:10.1016/s0896-6273(00)80658-3

5. Asaad WF, Rainer G, Miller EK. Task-Specific Neural Activity in the Primate Prefrontal Cortex. J Neurophysiol. 2000;84: 451–459. doi:10.1152/jn.2000.84.1.451

6. Genovesio A, Brasted PJ, Mitz AR, Wise SP. Prefrontal Cortex Activity Related to Abstract Response Strategies. Neuron. 2005;47: 307–320. doi:10.1016/j.neuron.2005.06.006

7. Petrides M, Pandya DN. Comparative cytoarchitectonic analysis of the human and the macaque ventrolateral prefrontal cortex and corticocortical connection patterns in the monkey. European Journal of Neuroscience. 2002;16: 291–310. doi:10.1046/j.1460-9568.2001.02090.x

8. Petrides M. Dorsolateral prefrontal cortex: Comparative cytoarchitectonic analysis”in the human and the macaque brain and corticocortical connection patterns. European Journal of Neuroscience. 1999;11: 1011– 1036. doi:10.1046/j.1460-9568.1999.00518.x

9. Borra E, Gerbella M, Rozzi S, Luppino G. Anatomical evidence for the involvement of the macaque ventrolateral prefrontal area 12r in controlling goal-directed actions. J Neurosci. 2011;31: 12351–63. doi:10.1523/JNEUROSCI.1745-11.2011

10. Gerbella M, Belmalih A, Borra E, Rozzi S, Luppino G. Multimodal architectonic subdivision of the caudal ventrolateral prefrontal cortex of the macaque monkey. Brain Struct Funct. 2007;212: 269–301. doi:10.1007/s00429-007-0158-9

11. Gerbella M, Borra E, Tonelli S, Rozzi S, Luppino G. Connectional heterogeneity of the ventral part of the macaque area 46. Cerebral Cortex. 2013;23: 967–987. doi:10.1093/cercor/bhs096

12. Barbas H. Anatomic organization of basoventral and mediodorsal visual recipient prefrontal regions in the rhesus monkey. Journal of Comparative Neurology. 1988;276: 313–342. doi:10.1002/cne.902760302

13. Barbas H, Mesulam MM. Organization of afferent input to subdivisions of area 8 in the rhesus monkey. J Comp Neurol. 1981;200: 407–31. doi:10.1002/cne.902000309

14. Barbas H, Mesulam MM. Cortical afferent input to the principalis region of the rhesus monkey. Neuroscience. 1985;15: 619–37. doi:10.1016/0306-4522(85)90064-8

15. Koechlin E, Ody C, Kouneiher F. The Architecture of Cognitive Control in the Human Prefrontal Cortex. Science (1979). 2003;302: 1181–1185. doi:10.1126/science.1088545

16. Nee DE, D’Esposito M. The hierarchical organization of the lateral prefrontal cortex. Elife. 2016;5. doi:10.7554/eLife.12112

17. Nee DE, D’Esposito M. Causal evidence for lateral prefrontal cortex dynamics supporting cognitive control. Elife. 2017;6. doi:10.7554/eLife.28040

18. Riley MR, Qi X-L, Constantinidis C. Functional specialization of areas along the anterior-posterior axis of the primate prefrontal cortex. Cereb Cortex. 2017;27: 3683–3697. doi:10.1093/cercor/bhw190

19. Riley MR, Qi XL, Zhou X, Constantinidis C. Anterior-posterior gradient of plasticity in primate prefrontal cortex. Nat Commun. 2018;9. doi:10.1038/s41467-018-06226-w

20. Tan PK, Tang C, Herikstad R, Pillay A, Libedinsky C. Distinct Lateral Prefrontal Regions Are Organized in an Anterior-Posterior Functional Gradient. J Neurosci. 2023;43: 6564–6572. doi:10.1523/JNEUROSCI.0007-23.2023

21. Abdallah M, Zanitti GE, Iovene V, Wassermann D. Functional gradients in the human lateral prefrontal cortex revealed by a comprehensive coordinate-based meta-analysis. Elife. 2022;11. doi:10.7554/eLife.76926

22. Goldman-Rakic PS. The prefrontal landscape: implications of functional architecture for understanding human mentation and the central executive. Philos Trans R Soc Lond B Biol Sci. 1996;351: 1445–53. doi:10.1098/rstb.1996.0129

23. Petrides M. Lateral prefrontal cortex: architectonic and functional organization. Philos Trans R Soc Lond B Biol Sci. 2005;360: 781–95. doi:10.1098/rstb.2005.1631

24. Koechlin E, Summerfield C. An information theoretical approach to prefrontal executive function. Trends Cogn Sci. 2007;11: 229–235. doi:10.1016/j.tics.2007.04.005

25. Badre D, Nee DE. Frontal Cortex and the Hierarchical Control of Behavior. Trends Cogn Sci. 2018;22: 170–188. doi:10.1016/j.tics.2017.11.005

26. Azuar C, Reyes P, Slachevsky A, Volle E, Kinkingnehun S, Kouneiher F, et al. Testing the model of caudo-rostral organization of cognitive control in the human with frontal lesions. Neuroimage. 2014;84: 1053–60. doi:10.1016/j.neuroimage.2013.09.031

27. Saleem KS, Miller B, Price JL. Subdivisions and connectional networks of the lateral prefrontal cortex in the macaque monkey. J Comp Neurol. 2014;522: 1641–90. doi:10.1002/cne.23498

28. Borra E, Gerbella M, Rozzi S, Luppino G. Projections from caudal ventrolateral prefrontal areas to brainstem preoculomotor structures and to Basal Ganglia and cerebellar oculomotor loops in the macaque. Cereb Cortex. 2015;25: 748–64. doi:10.1093/cercor/bht265

29. Gerbella M, Belmalih A, Borra E, Rozzi S, Luppino G. Cortical connections of the macaque caudal ventrolateral prefrontal areas 45A and 45B. Cereb Cortex. 2010;20: 141–68. doi:10.1093/cercor/bhp087

30. Borra E, Luppino G. Large-scale temporo–parieto–frontal networks for motor and cognitive motor functions in the primate brain. Cortex. Masson SpA; 2019. pp. 19–37. doi:10.1016/j.cortex.2018.09.024

31. Borra E, Gerbella M, Rozzi S, Luppino G. Anatomical evidence for the involvement of the macaque ventrolateral prefrontal area 12r in controlling goal-directed actions. Journal of Neuroscience. 2011;31: 12351–12363. doi:10.1523/JNEUROSCI.1745-11.2011

32. Borra E, Gerbella M, Rozzi S, Tonelli S, Luppino G. Projections to the superior colliculus from inferior parietal, ventral premotor, and ventrolateral prefrontal areas involved in controlling goal-directed hand actions in the macaque. Cereb Cortex. 2014;24: 1054–65. doi:10.1093/cercor/bhs392

33. Markowitz DA, Curtis CE, Pesaran B. Multiple component networks support working memory in prefrontal cortex. Proc Natl Acad Sci U S A. 2015;112: 11084–11089. doi:10.1073/pnas.1504172112

34. Trepka E, Spitmaan M, Qi X-L, Constantinidis C, Soltani A. Training-Dependent Gradients of Timescales of Neural Dynamics in the Primate Prefrontal Cortex and Their Contributions to Working Memory. J Neurosci. 2024;44. doi:10.1523/JNEUROSCI.2442-21.2023

35. Boch RA, Goldberg ME. Participation of prefrontal neurons in the preparation of visually guided eye movements in the rhesus monkey. J Neurophysiol. 1989;61: 1064–84. doi:10.1152/jn.1989.61.5.1064

36. Tanila H, Carlson S, Linnankoski I, Kahila H. Regional distribution of functions in dorsolateral prefrontal cortex of the the monkey. Behav Brain Res. 1993;53: 63–71. doi:10.1016/s0166-4328(05)80266-9

37. Hoshi E, Shima K, Tanji J. Task-Dependent Selectivity of Movement-Related Neuronal Activity in the Primate Prefrontal Cortex. J Neurophysiol. 1998;80: 3392–3397. doi:10.1152/jn.1998.80.6.3392

38. Simone L, Rozzi S, Bimbi M, Fogassi L. Movement-related activity during goal-directed hand actions in the monkey ventrolateral prefrontal cortex. Foxe J, editor. Eur J Neurosci. 2015;42: 2882–94. doi:10.1111/ejn.13040

39. Rozzi S, Fogassi L. Neural Coding for Action Execution and Action Observation in the Prefrontal Cortex and Its Role in the Organization of Socially Driven Behavior. Front Neurosci. 2017;11: 1–9. doi:10.3389/fnins.2017.00492

40. Gerbella M, Rozzi S, Rizzolatti G. The extended object-grasping network. Exp Brain Res. 2017;235: 2903– 2916. doi:10.1007/s00221-017-5007-3

41. Nelissen K, Borra E, Gerbella M, Rozzi S, Luppino G, Vanduffel W, et al. Action observation circuits in the macaque monkey cortex. J Neurosci. 2011;31: 3743–56. doi:10.1523/JNEUROSCI.4803-10.2011

42. Rozzi S, Bimbi M, Gravante A, Simone L, Fogassi L. Visual response of ventrolateral prefrontal neurons and their behavior-related modulation. Sci Rep. 2021;11: 10118. doi:10.1038/s41598-021-89500-0

43. Gerbella M, Borra E, Tonelli S, Rozzi S, Luppino G. Connectional Heterogeneity of the Ventral Part of the Macaque Area 46. Cerebral Cortex. 2013;23: 967–987. doi:10.1093/cercor/bhs096

44. Borra E, Gerbella M, Rozzi S, Luppino G. Anatomical evidence for the involvement of the macaque ventrolateral prefrontal area 12r in controlling goal-directed actions. Journal of Neuroscience. 2011;31: 12351–12363. doi:10.1523/JNEUROSCI.1745-11.2011

45. Gerbella M, Belmalih A, Borra E, Rozzi S, Luppino G. Cortical connections of the macaque caudal ventrolateral prefrontal areas 45A and 45B. Cereb Cortex. 2010;20: 141–168. doi:10.1093/cercor/bhp087

46. Nelissen K, Borra E, Gerbella M, Rozzi S, Luppino G, Vanduffel W, et al. Action observation circuits in the macaque monkey cortex. J Neurosci. 2011;31: 3743–56. doi:10.1523/JNEUROSCI.4803-10.2011

47. Rozzi S, Gravante A, Basile C, Cappellaro G, Gerbella M, Fogassi L. Ventrolateral prefrontal neurons of the monkey encode instructions in the ‘pragmatic’ format of the associated behavioral outcomes. Prog Neurobiol. 2023;229: 102499. doi:10.1016/j.pneurobio.2023.102499

48. Ó Scalaidhe SP, Wilson FAW, Goldman-Rakic PS. Areal Segregation of Face-Processing Neurons in Prefrontal Cortex. Science (1979). 1997;278: 1135–1138. doi:10.1126/science.278.5340.1135

49. Neal JW, Pearson RCA, Powell TPS. The ipsilateral corticocortical connections of area 7 with the frontal lobe in the monkey. Brain Res. 1990;509: 31–40. doi:10.1016/0006-8993(90)90305-U

50. Seltzer B, Pandya DN. Frontal lobe connections of the superior temporal sulcus in the rhesus monkey. J Comp Neurol. 1989;281: 97–113. doi:10.1002/cne.902810108

51. Cavada C, Goldman-Rakic PS. Posterior parietal cortex in rhesus monkey: II. Evidence for segregated corticocortical networks linking sensory and limbic areas with the frontal lobe. J Comp Neurol. 1989;287: 422–45. doi:10.1002/cne.902870403

52. Cavada C, Goldman-Rakic PS. Posterior parietal cortex in rhesus monkey: I. Parcellation of areas based on distinctive limbic and sensory corticocortical connections. J Comp Neurol. 1989;287: 393–421. doi:10.1002/cne.902870402

53. Petrides M, Pandya DN. Projections to the frontal cortex from the posterior parietal region in the rhesus monkey. J Comp Neurol. 1984;228: 105–16. doi:10.1002/cne.902280110

54. Chavis DA, Pandya DN. Further observations on corticofrontal connections in the rhesus monkey. Brain Res. 1976;117: 369–386. doi:10.1016/0006-8993(76)90089-5

55. Wilson FAW, Scalaidhe SPÓ, Goldman-Rakic PS. Dissociation of Object and Spatial Processing Domains in Primate Prefrontal Cortex. Science (1979). 1993;260: 1955–1958. doi:10.1126/science.8316836

56. Logothetis NK, Sheinberg DL. Visual Object Recognition. Annu Rev Neurosci. 1996;19: 577–621. doi:10.1146/annurev.ne.19.030196.003045

57. Tanaka K. Inferotemporal Cortex and Object Vision. Annu Rev Neurosci. 1996;19: 109–139. doi:10.1146/annurev.ne.19.030196.000545

58. Borra E, Luppino G. Comparative anatomy of the macaque and the human frontal oculomotor domain. Neurosci Biobehav Rev. 2021;126: 43–56. doi:10.1016/j.neubiorev.2021.03.013

59. Romanski LM. Representation and integration of auditory and visual stimuli in the primate ventral lateral prefrontal cortex. Cereb Cortex. 2007;17 Suppl 1: i61–9. doi:10.1093/cercor/bhm099

60. Romanski LM, Sharma KK. Multisensory interactions of face and vocal information during perception and memory in ventrolateral prefrontal cortex. Philos Trans R Soc Lond B Biol Sci. 2023;378: 20220343. doi:10.1098/rstb.2022.0343

61. Borra E, Gerbella M, Rozzi S, Luppino G. The macaque lateral grasping network: A neural substrate for generating purposeful hand actions. Neurosci Biobehav Rev. 2017;75: 65–90. doi:10.1016/j.neubiorev.2017.01.017

62. Wallis JD, Anderson KC, Miller EK. Single neurons in prefrontal cortex encode abstract rules. Nature. 2001;411: 953–956. doi:10.1038/35082081

63. Genovesio A, Brasted PJ, Wise SP. Representation of future and previous spatial goals by separate neural populations in prefrontal cortex. J Neurosci. 2006;26: 7305–16. doi:10.1523/JNEUROSCI.0699-06.2006

64. Tsujimoto S, Genovesio A, Wise SP. Transient neuronal correlations underlying goal selection and maintenance in prefrontal cortex. Cereb Cortex. 2008;18: 2748–61. doi:10.1093/cercor/bhn033

65. Rainer G, Rao SC, Miller EK. Prospective coding for objects in primate prefrontal cortex. J Neurosci. 1999;19: 5493–505. doi:10.1523/JNEUROSCI.19-13-05493.1999

66. Murata A, Gallese V, Luppino G, Kaseda M, Sakata H. Selectivity for the Shape, Size, and Orientation of Objects for Grasping in Neurons of Monkey Parietal Area AIP. J Neurophysiol. 2000;83: 2580–2601. doi:10.1152/jn.2000.83.5.2580

67. Sakata H, Taira M, Murata A, Mine S. Neural mechanisms of visual guidance of hand action in the parietal cortex of the monkey. Cereb Cortex. 1995;5: 429–38. doi:10.1093/cercor/5.5.429

68. Bichot NP, Heard MT, DeGennaro EM, Desimone R. A Source for Feature-Based Attention in the Prefrontal Cortex. Neuron. 2015;88: 832–844. doi:10.1016/j.neuron.2015.10.001

69. Wardak C, Vanduffel W, Orban GA. Searching for a salient target involves frontal regions. Cereb Cortex. 2010;20: 2464–77. doi:10.1093/cercor/bhp315

70. Soon CS, Brass M, Heinze HJ, Haynes JD. Unconscious determinants of free decisions in the human brain. Nat Neurosci. 2008;11: 543–545. doi:10.1038/nn.2112

71. Tsujimoto S, Genovesio A, Wise SP. Evaluating self-generated decisions in frontal pole cortex of monkeys. Nat Neurosci. 2010;13: 120–126. doi:10.1038/nn.2453

72. Gerbella M, Borra E, Tonelli S, Rozzi S, Luppino G. Connectional heterogeneity of the ventral part of the macaque area 46. Cereb Cortex. 2013;23: 967–87. doi:10.1093/cercor/bhs096

73. De Renzi E, Cavalleri F, Facchini S. Imitation and utilisation behaviour. J Neurol Neurosurg Psychiatry. 1996;61: 396–400. doi:10.1136/jnnp.61.4.396

74. Pacherie E. The anarchic hand syndrome and utilization behavior: a window onto agentive self-awareness. Funct Neurol. 2007;22: 211–7. Available: http://www.ncbi.nlm.nih.gov/pubmed/18182128

75. Lhermitte F. “Utilization behaviour” and its relation to lesions of the frontal lobes. Brain. 1983;106 (Pt 2): 237–55. doi:10.1093/brain/106.2.237

76. Rozzi S, Ferrari PF, Bonini L, Rizzolatti G, Fogassi L. Functional organization of inferior parietal lobule convexity in the macaque monkey: electrophysiological characterization of motor, sensory and mirror responses and their correlation with cytoarchitectonic areas. Eur J Neurosci. 2008;28: 1569–88. doi:10.1111/j.1460-9568.2008.06395.x

77. Kobak D, Brendel W, Constantinidis C, Feierstein CE, Kepecs A, Mainen ZF, et al. Demixed principal component analysis of neural population data. Elife. 2016;5. doi:10.7554/eLife.10989

78. Meyers EM. The neural decoding toolbox. Front Neuroinform. 2013;7. doi:10.3389/fninf.2013.00008

79. Ceccarelli F, Ferrucci L, Londei F, Ramawat S, Brunamonti E, Genovesio A. Static and dynamic coding in distinct cell types during associative learning in the prefrontal cortex. Nat Commun. 2023;14: 8325. doi:10.1038/s41467-023-43712-2

80. Spaak E, Watanabe K, Funahashi S, Stokes MG. Stable and Dynamic Coding for Working Memory in Primate Prefrontal Cortex. J Neurosci. 2017;37: 6503–6516. doi:10.1523/JNEUROSCI.3364-16.2017

81. Maris E, Oostenveld R. Nonparametric statistical testing of EEG- and MEG-data. J Neurosci Methods. 2007;164: 177–190. doi:10.1016/j.jneumeth.2007.03.024

